# Production of β-PheRS fragments correlates with food avoidance and slow growth, and is suppressed by the appetite-inducing hormone CCHa2

**DOI:** 10.1101/2023.01.11.523627

**Authors:** Dominique Brunßen, Beat Suter

## Abstract

The housekeeping tRNA synthetases play many non-canonical roles with diverse functions. The phenylalanyl-tRNA synthetase (PheRS/FARS) is an α_2_β_2_ tetramere. Recently, human patients with mutations in *FARSB*, the homolog of *β-PheRS* in Drosophila, have been reported to display problems gaining weight. Here, we show in Drosophila that overexpressing the β subunit in the context of the complete PheRS leads to larval roaming, food avoidance, slow growth, and a developmental delay that can last several days and even prevents pupation. Narrowing down the tissue involved in this behavioral and developmental effect revealed that expression in CCHa2^+^ and Pros^+^ cells induced this phenotype. Simultaneous expression of β-PheRS, α-PheRS, and the appetite-inducing CCHa2 peptide rescued these phenotypes, linking this *β-PheRS* activity to the appetite-controlling pathway. The fragmentation dynamics of the excessive β-PheRS points to a β-PheRS fragment as a likely candidate inducer of these phenotypes. Fragmentation of PheRS (FARS) has also been observed in humans and mutations in human *β-PheRS (FARSB)* can lead to problems in gaining weight. This study, therefore, points to a potential mechanism for the human phenotype and to possible novel approaches to research ways to correct the balance between hunger and satiety signals in the context of obesity.

## Introduction

In *Drosophila melanogaster*, charging of all tRNAs is performed by 34 aaRSs (Lu et al 2015). Whereas 15 aaRSs are exclusively cytoplasmic, 15 are mitochondrial and 4 are functioning in the cytoplasm and the mitochondria. aaRSs function primarily in loading the appropriate tRNA with its cognate amino acid. However, over the last couple of decades, many additional functions, so-called non-canonical functions, were discovered for multiple tRNA synthetases. The non-canonical functions of aaRSs range from extracellular functions, such as cytokines or regulators of angiogenesis, to retroviral particle assembly and to cytoplasmic functions as regulators of apoptosis and translation. Involvements in DNA replication or synthesis, 3’-end formation of mRNAs, tRNA export from the nucleus, and import into mitochondria have been described, too (Smirnova et al 2012, Park et al 2005 a). Furthermore, the human tyrosyl-tRNA synthetase (TyrRS) can be secreted to function in the extracellular space as a signal after it is split into two polypeptide fragments that act as cytokines with two different activities (Wakasugi and Schimmel 1999, Wakasugi et al 2002). Also, human tryptophanyl-tRNA synthetase (TrpRS) is secreted and performs an extracellular, antiangiogenic activity (Wakasugi et al 2002). And human lysyl-tRNA synthetase (LysRS) is secreted upon TNF-α activation and triggers a proinflammatory response (Park et al 2005 b). An example of a cytoplasmic non-canonical function is provided by the glutamyl-prolyl-tRNA synthetase (GluProRS) which can cause gene-specific translational silencing (Sampath et al 2004). Further functions were also shown for KRS, phenylalanyl-tRNA synthetase (PheRS), threonyl-tRNA synthetase (ThrRS), and seryl-tRNA synthetase (SerRS) which are associated with the activation of DNA replication while glutaminyl-tRNA synthetase (GlnRS), and GluProRS are associated with inflammation and apoptosis (Jung 2015). GluProRS is additionally found as a mTORC1-S6K1 target that is involved in the regulation of adiposity and aging in mice (Arif et al 2017).

The cytoplasmic PheRS/FARS is a large and complex tRNA synthetase with a heterotetrameric structure consisting of two α and two β subunits. PheRS is markedly conserved in all species throughout evolution (Finarov et al 2010). The α-PheRS with the active site and the β-PheRS with the tRNA^Phe^ recognition site are only functional in aminoacylation in the complex (Finarov et al 2010). In most cells, the two PheRS subunits stabilize each other (Lu et al 2014). Lower levels of one subunit result in a decrease in the other subunit (Lu et al 2014, Xu et al 2018) whereas increasing levels of β-PheRS require co-overexpression of α-PheRS (Lu et al 2014, Ho et al 2021).

Recently, Ho et al (2021) discovered an aminoacylation- and translation-independent function of *Drosophila* α-PheRS. An active site mutant form of α-PheRS can accelerate growth and proliferation. Furthermore, an α-PheRS fragment that is present in some tissues counteracts Notch signaling in situations where this signal affects tissue homeostasis by either promoting stem cell fate or differentiation (Ho et al, 2022).

Among the few reports describing possible non-canonical functions of PheRS is a study performed in rats. *β-PheRS (FARSB), isoleucyl-tRNA synthetase (IleRS)*, and *methionyl-tRNA synthetase (MetRS)* mRNA expression levels increase in spinal dorsal horn neurons upon peripheral nerve injury (Park et al 2015). This suggested the possibility that FARSB, IARS, and MARS may act as neurotransmitters for transferring abnormal sensory signals after peripheral nerve damage (Park et al 2015).

The B5 domain of β-PheRS is not involved in aminoacylation but conserved through evolution, suggesting that it might provide a different, non-canonical, activity. Human B5 contains a “helix-turn-helix” motive and two B5 domains hB5 and hB5*. With their particular distance from each other, these can bind DNA, forming a loop between them. The target DNA does not need to contain a specified motive but a specified length of 80 base pairs (Dou et al, 2001). A related function as an mRNA binding protein was also suggested for PheRS after it was found in a screen as a possible mRNA binding protein (Castello et al. 2012).

Recently the first human patients with mutations in FARSB were described. Trans-heterozygous (also called compound heterozygous) mutations in FARSB were viable with severe health and growth problems (Xu et al 2018, Antonellis et al 2018, Zadjali et al 2018). The FARSB mutations caused growth restriction, brain calcification, and interstitial lung disease. The mutations in FARSB can also lead to lower protein levels of FARSB and its partner alpha subunit (Xu et al 2018, Antonellis et al 2018, Zadjali et al 2018), but this did not seem to affect translation, suggesting that a non-canonical effect caused the clinical condition through an unknown mechanism (Xu et al 2018).

In this study, we show that manipulating Drosophila PheRS/FARS levels and subunit expression can affect growth speed, the timing of pupation, and also behavioral aspects, such as feeding and roaming. We also present evidence for the involvement of specific brain cells and/or intestinal cells in this process. We further discuss the evidence that a cleavage product of β-PheRS provides the activity for this process.

## Materials and Methods

### Material

#### Key resource table

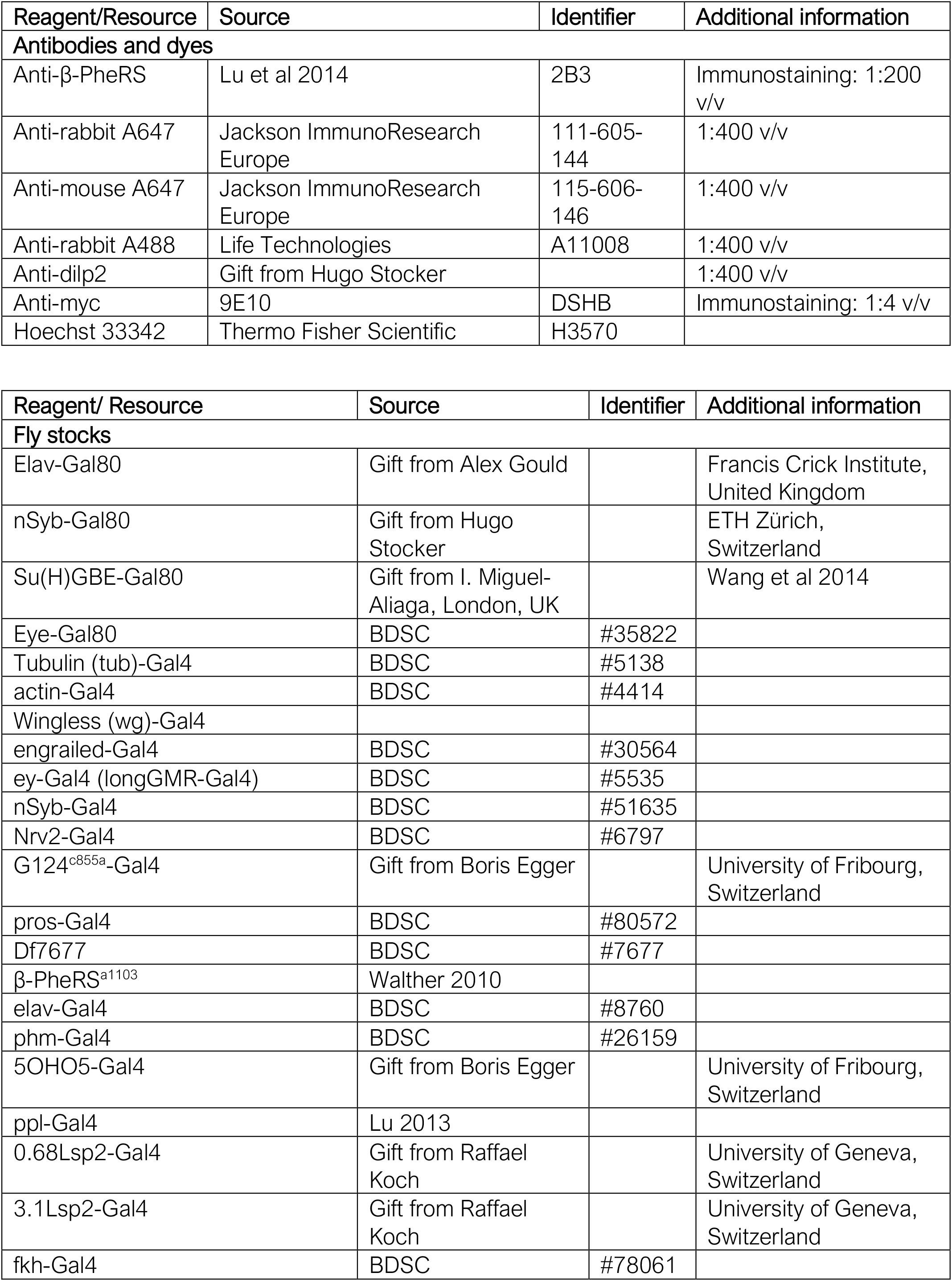
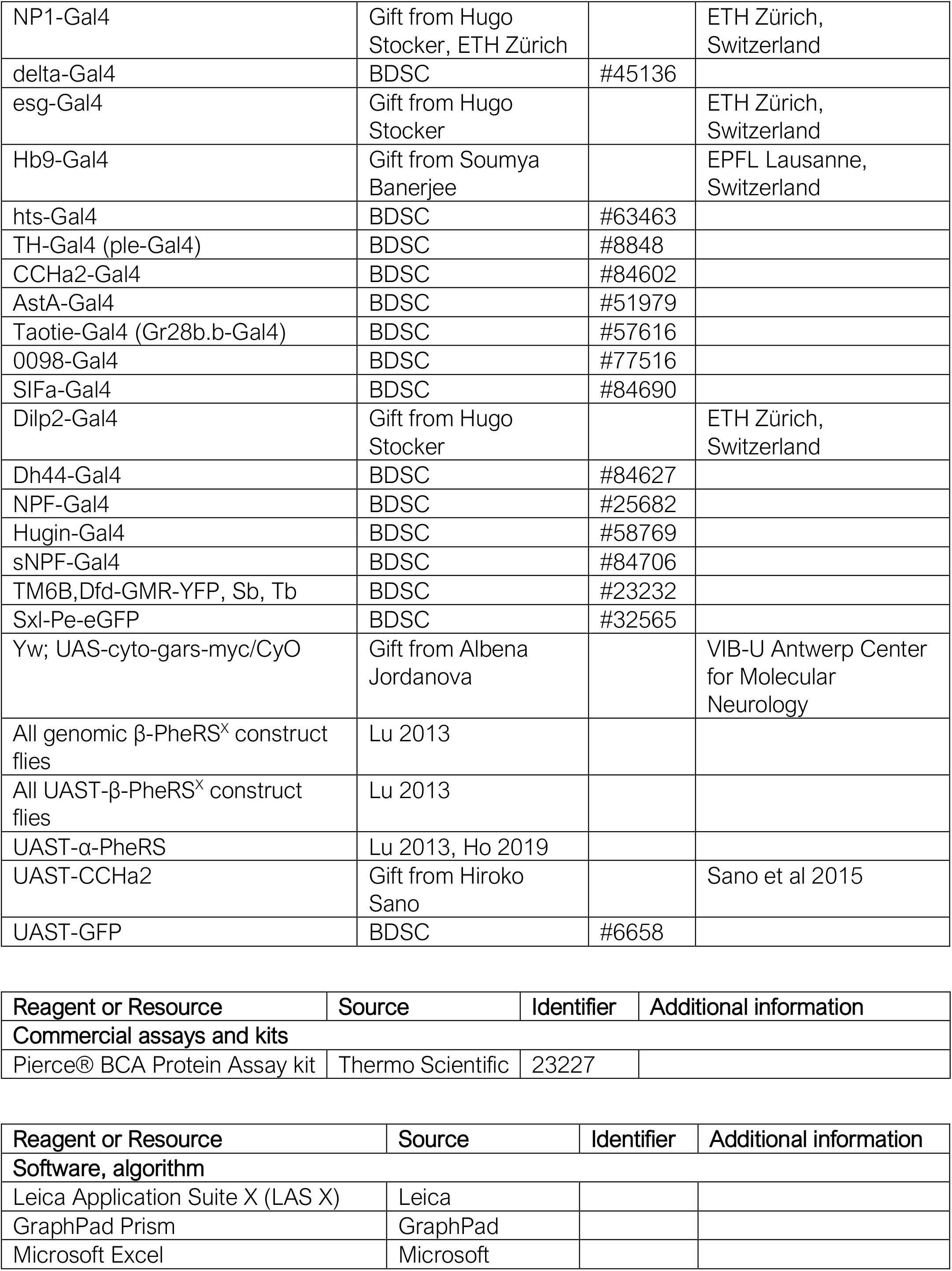

#### Buffers

|  |  |  |
| --- | --- | --- |
| **Fly food recipe**<br>20.4 l H<sub>2</sub>O<br>1.68 g Maize flour<br>720 g Yeast<br>1.8 g Syrup | **Apple juice plates**<br>1 l H<sub>2</sub>O tap<br>30 g Agar<br>350 ml Apple juice<br>35 g sugar | **10X PBS**<br>40 g NaCl,<br>1 g KCl,<br>7.2 g Na<sub>2</sub>HPO<sub>4</sub>,<br>1.2 g KH<sub>2</sub>PO<sub>4</sub>, |
| 192 g potassium sodium tartrate tetrahydrate<br>36 g Nipagin<br>120 ml propionic acid | 2 g nipagin | pH adjusted to 7.4<br>filled up to 0.5 L |
| **PFA-fix**<br>1X PBSTT<br>4% Paraformaldehyde (w/v) | **PBST**<br>1X PBS 10X<br>0.2 % Tween-20 | **PBSTT**<br>1X PBST<br>0.1 % TritonX-100 |
| **Staining blocking solution**<br>1X PBSTT<br>5% non-fat dry milk (w/v) |  |  |

## Methods

### Fly keeping

Stocks in use were kept at 25°C in glass vials or plastic bottles with a day/night cycle (12h/12h). Larval experiments were performed at 25°C in a 24h dark incubator. Stocks for long-term keeping were kept at 18°C in glass vials with a day/night (12h/12h) cycle on standard food.

### Time to pupation assay and lethality

Egg lays were performed on day 0 between 9 am -1 pm on apple juice plates with yeast paste. 50 L1 larvae not displaying the Dfd-GMR-YFP signal from the balancer were collected on day 1 between 2 - 4 pm and placed onto new apple juice plates with yeast paste. The larvae were kept at 25°C in 24h darkness. From day 4 on, every day at 5 pm the pupae were counted. The mean time to pupation was calculated with GraphPad Prism. The lethal larvae were determined as follows: collected larvae (50), minus the number of pupae, which were counted at the end of the pupation assay. All results represent biological duplicates or triplicates.

### Immunofluorescent staining and confocal microscopy

The tissue of interest was dissected in *PBS* (max 30 min) and fixed with *PFA-fix*. Wing discs and brains were fixed for 40 min and guts for 1h. Fixed tissue was rinsed 3x and washed 3x 10 min with *PBSTT* and blocked with *staining blocking solution* for 2h at room temperature. The 1^st^ Ab was added to the *staining blocking solution* overnight at 4°C, rinsed 3x, and washed 3x for 20 min. Secondary antibodies were added in *staining blocking solution* for 4 hours at room temperature, rinsed 3x, and washed for 20 min. Hoechst 33258 (5 µg/ml in PBST) was added for 20 min. The tissue was then washed again 2x for 20 min and mounted with Aqua/Poly Mount (Polysciences Inc., US). Image acquisition was performed on a Leica SP8 confocal microscope. The recipes of all solutions are noted in the *Key Resource Table*.

### Roaming Assessment

Egg lays were performed on day 0 between 10-12 am on apple juice plates with yeast paste. 50 L1 larvae not displaying the Dfd-GMR-YFP signal from the balancer were collected on day 1 between 2 – 4 pm and placed onto new apple juice plates with yeast paste. The larvae were kept at 25°C in 24h darkness. On day 4 at 9 am, a picture was taken, and the roaming larvae were counted. All results represent biological duplicates.

### Larval weight development and pupal weight measurement

Egg lays were prepared on day 0 between 10-12 am on apple juice plates with yeast paste. 4 hours later, the eggs with a Sxl-Pe-eGFP fluorescence signal were collected (female eggs were selected). 50 L1 larvae not displaying the Dfd-GMR-YFP signal from the balancer were collected on day 1 between 2 – 4 pm and put onto new apple juice plates with yeast paste. The larvae were kept at 25°C in 24h darkness. From day 3 on, the larval weight was measured individually. The pupal weight was measured within the first 24h after pupation. Statistical analysis was performed with GraphPad Prism.

### Measuring time to adulthood

Crosses were performed on standard food and the parental flies were transferred to new vials after two days. Two days later, they were removed from the second vials. Eclosed flies were sorted and counted every day for three days. When flies containing the Gal4 driver and flies containing the balancer eclosed from the first day on, the Gal4 driver was considered to not prolong the L3 phase.

### MS analysis

Egg laying was performed on apple juice plates without yeast for 2 hours. 26 hours later, 200 L1 larvae were collected without yeast contamination. The larvae were smashed in 100 µl urea buffer provided by the MS facility. Protein concentration was measured with the Pierce® BCA kit (Thermo Scientific) and the samples were analyzed by the MS facility. Experiments were performed in biological triplicates.

Mass Spectrometry analysis by the MS facility: Smashed larvae were reduced, alkylated, and precipitated overnight at -20°C with 5 volumes of acetone. The pellet was re-suspended in 8M urea/50mM Tris/HCl pH 8.00 to a protein concentration of 1mg/mL. Aliquots of 10µg protein were double digested with LysC (Promega, ratio 1:100) for 2hours at 37°C followed by Trypsin (Promega, ratio 1:100) at room temperature overnight. Digests were analysed in random order by loading 500ng onto a pre-column (C18 PepMap 100, 5µm, 100A, 300µm i.d. x 5mm length) at a flow rate of 50µL/min with solvent C (0.05% TFA in water/acetonitrile 98:2). After loading, peptides were eluted in back flush mode onto a home packed analytical Nano-column (Reprosil Pur C18-AQ, 1.9µm, 120A, 0.075 mm i.d. x 300mm length) using an acetonitrile gradient of 5% to 40% solvent B (0.1% Formic Acid in water/acetonitrile 4,9:95) in 180min at a flow rate of 250nL/min. The column effluent was directly coupled to a Fusion LUMOS mass spectrometer (Thermo Fischer, Bremen; Germany) via a nano-spray ESI source. Data acquisition was made in data dependent mode with precursor ion scans recorded in the orbitrap at resolution of 120’000 (at m/z=250) parallel to top speed fragment spectra of the most intense precursor ions in the linear trap for a cycle time of 3 seconds maximum. The HCD fragmentation type was applied for charge states 2 and 3, and ETD fragmentation for 4 to 9.

The mass spectrometry data was searched and quantified with MaxQuant (Cox et al 2008) (version 1.5.4.1) using the Drosophila melanogaster uniprot (uniport) database (release August 2017), to which common contaminants where added. The following parameters were used: digestion set to Trypsin/P, with maximum 3 missed cleavages; first search peptide tolerance set to 10 ppm, and MS/MS match tolerance to 0.4 Daltons. Carbamidomethylation on cysteine was given as a fixed modification; variables modifications were methionine oxidation, phenylalanylation and protein N-terminal acetylation. Match between runs was not enabled. Protein intensities were reported as MaxQuant’s Label Free Quantification (LFQ) values. Imputation and comparisons were performed for those protein groups for which there were at least 2 identifications in at least 1 group of replicates; re-normalisation, filtering and imputation were done with the DEP R package (Zhang et al 2018), using variance stabilisation normalisation (Huber et al 2002) and “MinProb” imputation method (draws from the 0.01th quantile). Differential expression tests were performed using empirical Bayes (moderated t-test) implemented in the R limma package (Kammers et al 2015). The Benjamini and Hochberg (Benjamini et al 1995) method was further applied to correct for multiple testing.

## Results

Two motives in the β-PheRS B5 domain are essential for β-PheRS stability and function The PheRS subunits can be divided into domains fulfilling different functions. While the functions of most domains are known, the biological function of the B5 domain of the β subunit (B5) is still unknown. B5 is not directly involved in aminoacylation but binds DNA and possibly also mRNA (Dou et al, 2001; Castello et al. 2012). In an attempt to investigate the function of B5, the codons for 5 potentially crucial β-PheRS residues and motives were mutated. These sites were selected because of their conservation and because they might be important for DNA or RNA binding (Fig. 1, Table 1, Dou et al 2001, Lu 2013). Mutant *β-PheRS* transgenes under their native promoter were then tested in the *β-PheRS*^*null*^ background. Three of the five mutants fully rescued the mutant phenotype, indicating that they still provided sufficient activity to perform the essential canonical function of *β-PheRS*. These three mutants were not further analyzed. The remaining two, *β-PheRS*^*B5a*^ and *β-PheRS*^*B5b*^ did not rescue, indicating that they were not functional (Table 1). This points to the R^353^ residue and the GYNN^371-4^ motive as essential for viability. Interestingly, these two sites are the most conserved ones (Fig. 1). Larvae containing either *β-PheRS*^*B5a*^ or *β-PheRS*^*B5b*^ in the *β-PheRS*^*null*^ background showed the same phenotype as the *β-PheRS*^*null*^ larvae. They hatched from the egg and appeared healthy. First instar larvae (L1 larvae) initially moved normally, but they did not grow and died during the first instar, a few hours after hatching. This is a common observation for essential genes in Drosophila, where the maternal contribution of most gene products allows development of embryos into larvae.

**Table 1:** Mutated residues & motives and the rescue ability of the mutated constructs

| Allele | Mutated Residues/Motives | Mutated to | Rescues β-PheRS<sup>null</sup> |
| --- | --- | --- | --- |
| B5a | R<sup>353</sup> | A | No |
| B5b | GYNN<sup>371-4</sup> | AAAA | No |
| B5c | DGDS<sup>341-4</sup> | AAAA | Yes |
| B5d | H<sup>358</sup> | A | Yes |
| B5e | KK<sup>376-7</sup> | AA | Yes |

**Figure 1:**
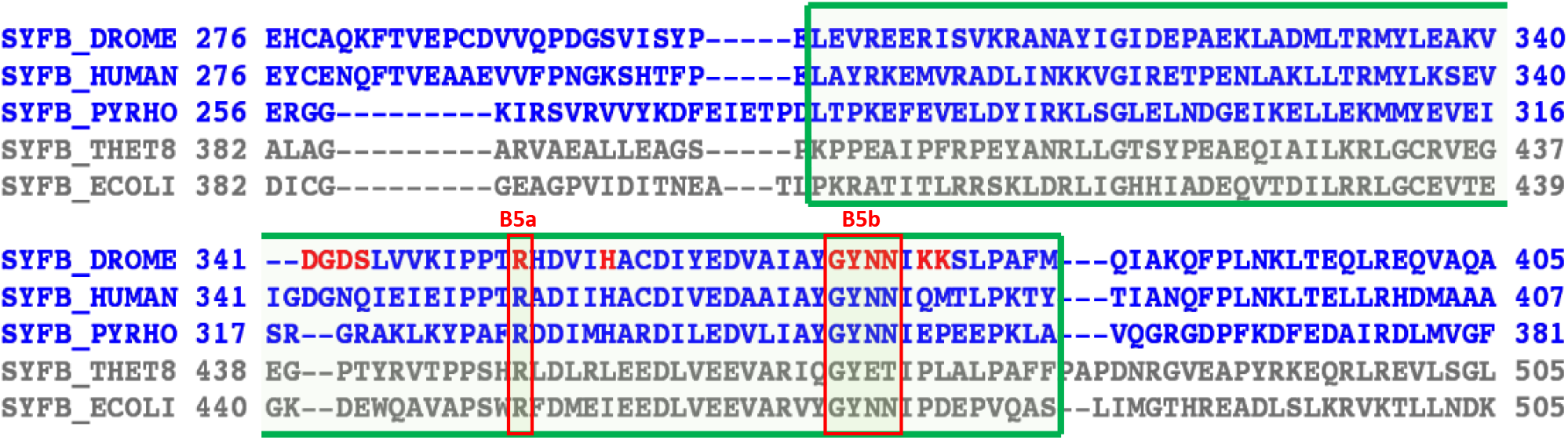
Structure based alignment of β-PheRS B5 domain and adjacent regions using the PROMALS3D method. The sequences of prokaryotes and archaea/eukaryotes are shown in gray and blue, respectively. Amino acid residues labeled in red are predicted to bind DNA/ RNA (Dou et al 2001; Wang and Brown 2006). DROME, *Drosophila* melanogaster, Human, homo sapiens; PYRHO, P.horikoshii; THET8, T thermophilus; ECOLI, E. coli. from Lu 2013). Marked with the green box is the B5 domain. Marked with the red box are the mutated sites B5a and B5b. The residue R^353^ and the motive GYNN^371-4^ were replaced by Alanins.

### Strong overexpression of α-/β-PheRS^+^, α-/β-PheRS^B5a^ or α-/β-PheRS^B5b^ leads to a developmental delay

An overexpression approach was used next. *α-PheRS, β-*PheRS^+^, *β-PheRS*^*B5a*^, *and β-PheRS*^*B5b*^ cDNAs were cloned behind the UAST promoter and overexpressed in the *α-/β-PheRS*^*+*^ background. Overexpressing β-PheRS^+^, β-PheRS^B5a^, and β-PheRS^B5b^ alone with the *actin-Gal4* or *tubulin-Gal4* (*tub-Gal4*) drivers did not lead to any phenotype. Similarly, no phenotype was seen when overexpressing β-PheRS^+^, β-PheRS^B5a^, and β-PheRS^B5b^ together with α-PheRS using the *actin-Gal4* driver. Interestingly, however, overexpression of the α-/β-PheRS^+^ subunits simultaneously with the strong *tub-Gal4* driver led to a slight developmental delay of 0-3 days. This phenotype became much more pronounced when overexpressing the α- and the mutant β-PheRS^B5a^ or β-PheRS^B5b^ subunits simultaneously. In this case, a developmental delay of 2-12 days for β-PheRS^B5a^ and 0-11 days for β-PheRS^B5b^ was observed (Table 2).

**Table 2:**
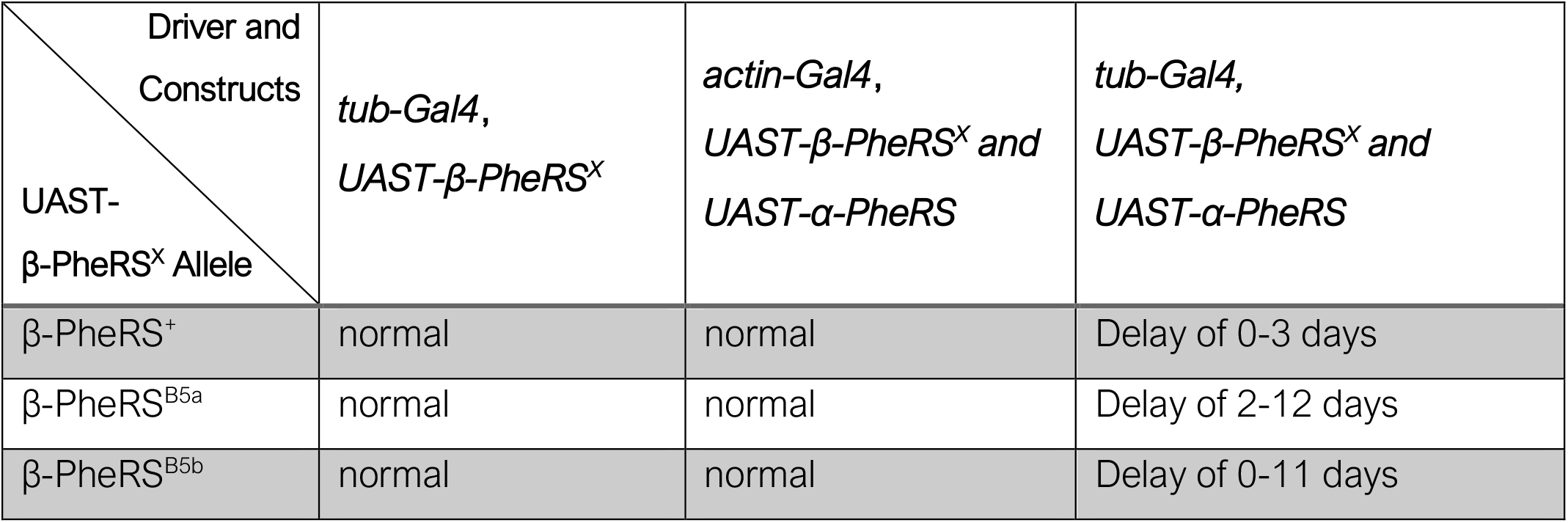
Time to pupation upon expression of α- and/or β-PheRS^X^ in the *α-/β-PheRS*^*+*^ background

The time from egg lay till the larvae started to pupate (time to pupation) was determined. It took the control larvae 5 days till the median of the larvae had pupated. Upon overexpression of α-/β-PheRS^+^, α-/β-PheRS^B5b^, and α-/β-PheRS^B5a^, respectively, it took the larvae 6 days, 9 days, and 10 days, respectively (Fig. 2A, Table 3).

**Table 3:** Median time till larvae pupated with tub-Gal4 overexpression and X-Gal80 inhibition

| Driver and inhibitor<br>Constructs | *tub-Gal4* | *tub-Gal4* and *elav-Gal80* | *tub-Gal4* and *nSyb-Gal80* | *tub-Gal4* and *Su(H)GBE-Gal80* | *tub-Gal4* and *eye-Gal80* |
| --- | --- | --- | --- | --- | --- |
| GFP o/e | 5 | 5 | 5 | 5 | 5 |
| α- and β-PheRS<sup>+</sup> o/e | 6 | 5 | 5 | 5 | 6 |
| α- and β-PheRS<sup>B5b</sup> o/e | 9 | 5 | 6 | 7 | 9 |
| α- and β-PheRS<sup>B5a</sup> o/e | 10 | 5 | 6 | 7 | 9 |

**Figure 2:**
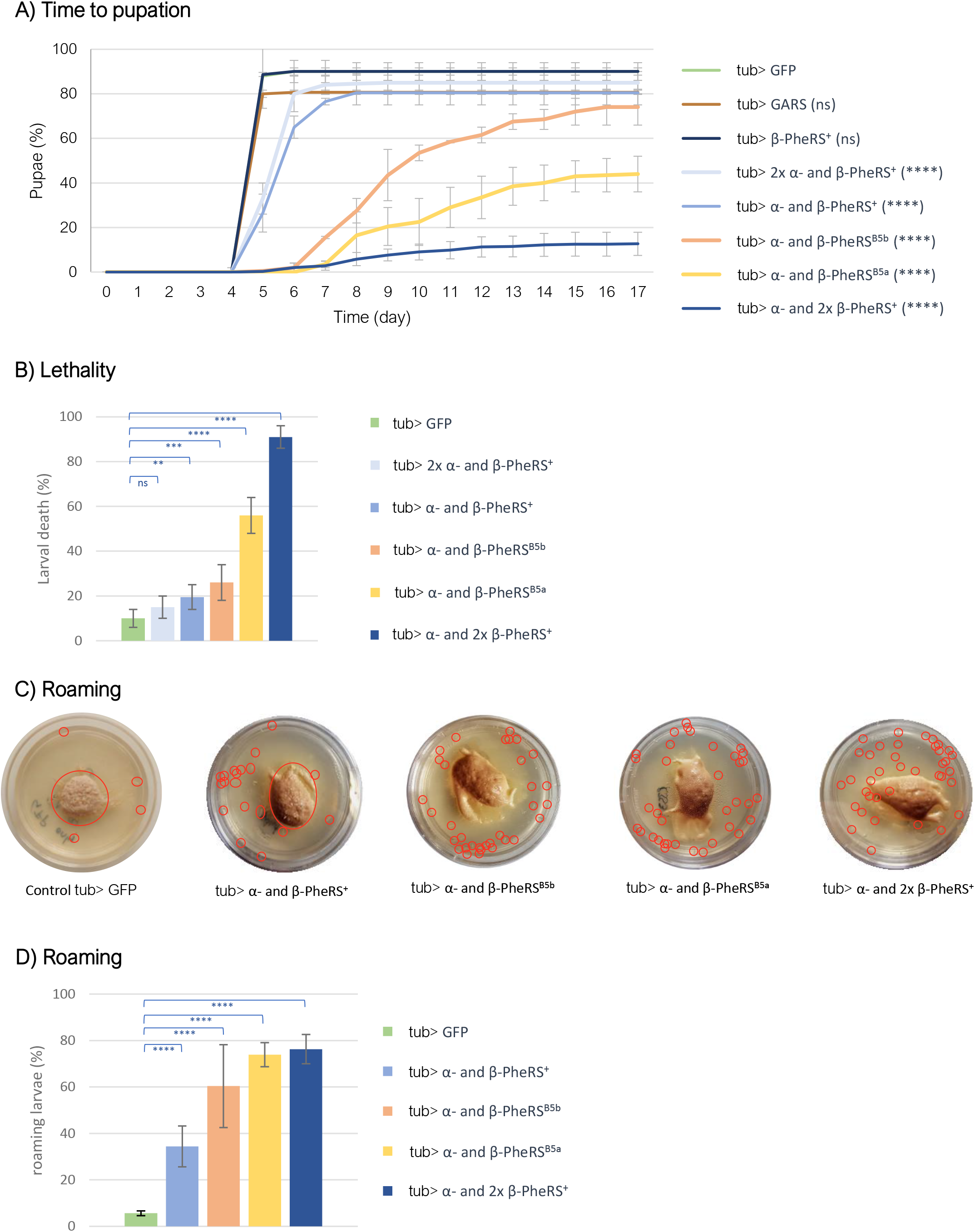
Effects of overexpression of PheRS subunits. (A) Time to pupation for the control, α-/β-PheRS^+^, α-/β-PheRS^B5a^ and α-/β-PheRS^B5b^ overexpressing larvae. One copy of the UAST driven transgenes was used for each subunit unless indicated otherwise. 2x means two copies of the transgene were tested (homozygous transgene) (Mann-Whitney-U-Test). (B) Lethality caused by overexpression of α-/β-PheRS (Fisher’s exact test). (C) L3 larvae overexpressing GFP or different PheRS subunit combinations were tested for their effect on feeding, roaming and survival on apple juice plates supplemented with yeast. The larvae are marked with red circles. (D) The percentage of roaming larvae was calculated (Fisher’s exact test). All experiments were performed in duplicates with each 50-100 larvae. Graphs represent median ± SD. Indicated statistical tests were used to compare results to control. p-value not significant (ns) > 0.05, * ≤ 0.05, ** ≤ 0.01, *** ≤ 0.001, **** ≤0.0001.

Interestingly, overexpression of α-/β-PheRS^+^ with an additional *β-PheRS*^*+*^ construct (one α-construct and 2 *β-PheRS*^*+*^ constructs) induced 76 % of the larvae to roam at day 4 and 91 % of the collected L1 larvae died as larvae (Fig 2B and 2C). Roaming larvae wandered away from the food and many kept roaming and probably died due to starvation, but a fraction of them even crawled out of the dish and dried out (Fig. 2B). Only 9 % of the collected and counted L1 larvae reached the pupal stage within 5 to 12 days after egg lay (Fig. 2A). In contrast, overexpression of α-/β-PheRS^+^ with an additional *α-PheRS*^*+*^ construct induced a slightly milder developmental delay compared to overexpression of one copy each of α-/β-PheRS^+^ (Fig 2A). This indicates that β-PheRS and not α-PheRS is the main factor inducing the developmental delay.

Overexpressing α-/β-PheRS^B5a^ or α-/β-PheRS^B5b^ produced a stronger phenotype than the α-/β-PheRS^+^ overexpression but a less severe phenotype than the 1x α- and 2x β-PheRS^+^ overexpression. This might indicate that the B5 mutations make β-PheRS more active for this secondary activity and possibly less active for its canonical function.

### Overexpression of α-/β-PheRS induces roaming

*Drosophila* grows exclusively during the larval stages. During this short 4-day period, the larvae grow from approximately 0.01 mg to approximately 2 mg, corresponding to a body mass increase of ∼200-fold (Tennessen and Thummel 2011). To attain this fast growth, the *Drosophila* larvae are feeding as much as possible during all larval stages and they only stop feeding once they have reached the optimal weight for pupation. Food, rich in protein, lipids, and other factors is required to support this rapid growth. Yeast is usually used as the rich fly food medium because it provides these ingrediencies. Once the larvae have reached their pupation size, they start to wander away from the food during the so-called wandering stage and pupate (Miroschnikow et al 2020).

To study this growth phase, we grew larvae on apple juice plates supplemented with yeast paste in the center of the plate. Control larvae, which over-expressed GFP, were mostly seen feeding on the yeast paste (Fig. 2C) till the late L3 stage, just before they entered the wandering L3 stage. In contrast, larvae overexpressing one copy each of α-/β-PheRS^+^, α-/β-PheRS^B5a^, and α-/β-PheRS^B5b^, respectively, tended to roam during all larval stages (Fig. 2C). Whereas only 6% of the control larvae were roaming at day 4, during the early L3 phase, upon overexpression of α-/β-PheRS^+^, α-/β-PheRS^B5a^, and α-/β-PheRS^B5b^, respectively, 34 %, 60 %, and 74 %, respectively, were observed roaming (Fig. 2D).

### Overexpression of α-/β-PheRS decreases growth speed

Developmental delay can be caused through different mechanisms. One is a failure in the initiation of pupation. In this case, the larval growth speed remains normal. This phenotype is seen in mutants affecting the Prothoracicotropic hormone (PTTH) and the Ecdysone pathway (Siegmund and Korge 2001, Quinn et al 2012, Thummel 2001). Loss of *Torso* or *PTTH* results in lower Ecdysone levels and this delays pupation due to a failure in the initiation of pupation. Normal growth speed combined with delayed initiation of pupation leads to larval overgrowth and enlarged pupae (Rewitz et al 2009). In contrast, *KDM5* is a gene encoding a transcription regulator which is essential for the prothoracic gland function. *KDM5* mutants show reduced growth rates, and this leads to a delay in pupation (Drelon et al 2019). To distinguish between these two possible mechanisms leading to a delay in pupation, the growth speed upon α-/β-PheRS^+^, α-/β-PheRS^B5a^ or α-/β-PheRS^B5b^ (α-/β-PheRS^X^) overexpression was measured. The weight of individual larvae was monitored. Even though we restricted the monitoring to the growth of female larvae, the time to pupation of the animals overexpressing α-/β-PheRS^B5a^ or α-/β-PheRS^B5b^ was very variable even between animals of the same genotype. Overexpression of glycyl-tRNA synthetase (GARS) was used as a control. Overexpression of α-/β-PheRS^+^ led to a small delay in growth and a difference in the max weight of 0.2 mg (7 %) and their average pupal weight was 0.2 mg (11 %) lighter (Fig. 3), whereas overexpression of α-/β-PheRS^B5a^ or α-/β-PheRS^B5b^ led to a much more striking delay in growth. On average, these larvae reached their maximal weight 149h and 174h, respectively, after egg lay with a difference of 0.7 mg (32 %) and 0.9 mg (44 %), respectively, and 0.6 mg (35 % and 36 %) lighter pupae compared to the control (Fig. 3). The growth is even more impaired in larvae overexpressing 1x α-and 2x β-PheRS^+^ (Suppl. Fig. S1). Most of these larvae died as L3 larvae after they had stopped growing (Fig. 2B).

**Figure 3:**
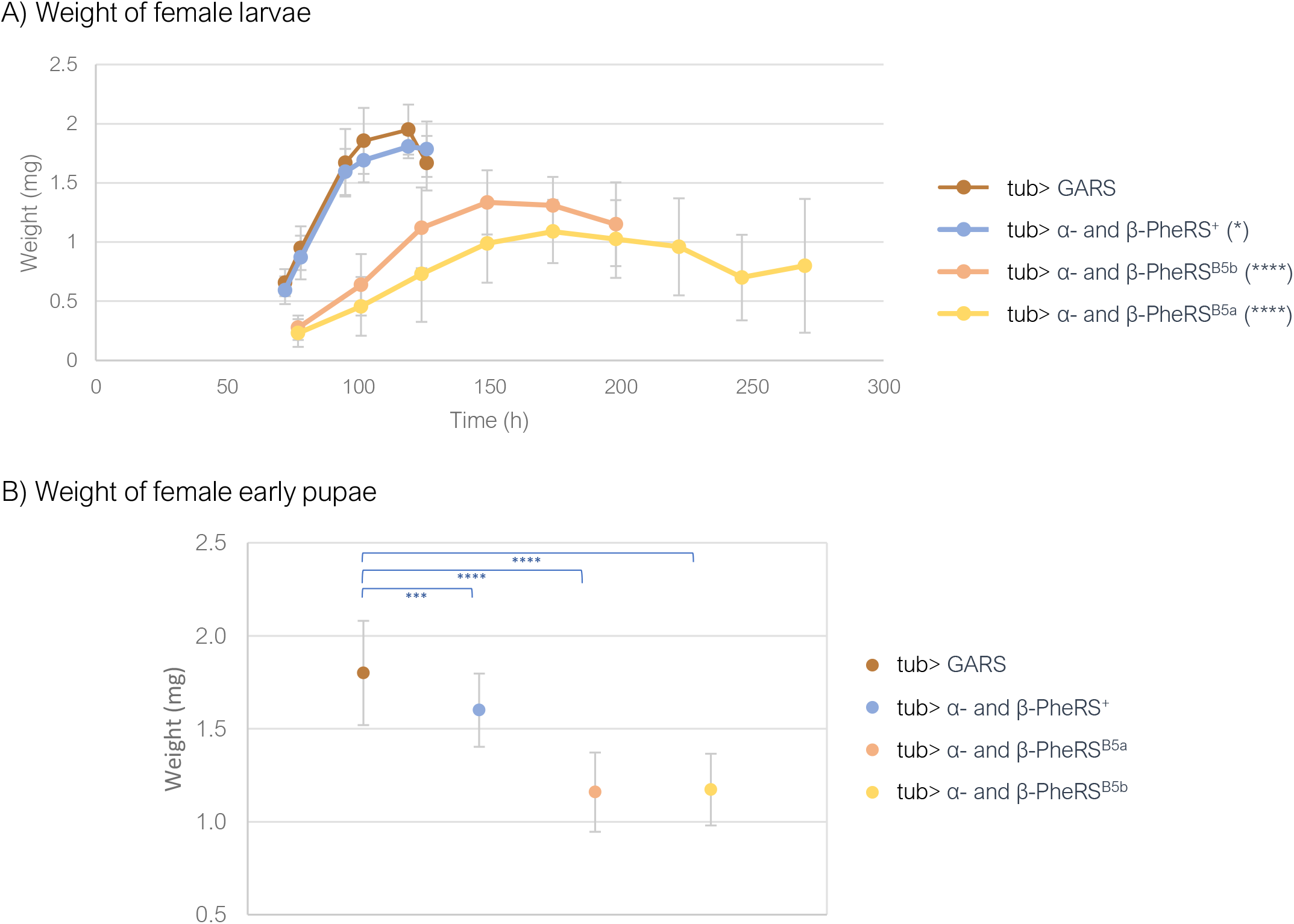
Weight of larvae and early pupae overexpressing PheRS subunits. (A) Weight development of control (tub> GARS) female larvae and female larvae overexpressing α-/β-PheRS^X^. Measurements were started on day 3 after egg-lay (Mann-Whitney-U-Test). (B) Pupal weight of control GARS overexpressing animals and α-/β-PheRS^X^ overexpressing animals was measured (One-way ANOVA). Graphs represent median ± SD, n = 50. Indicated statistical tests were used to compare results to control. p-value not significant (ns) > 0.05, * ≤ 0.05, ** ≤ 0.01, *** ≤ 0.001, **** ≤0.0001.

### Upon ubiquitous overexpression of PheRS, β-PheRS accumulates in IPCs

Upon overexpression of α-/β-PheRS with the ubiquitous driver *tub-Gal4*, the β-PheRS staining signal accumulates at higher levels only in a subset of tissues and cells while others show the normal signal levels, pointing to an active and tight control mechanism that restricts cellular PheRS levels in most cells. Cells that do not implement the β-PheRS level control are the segmentally organized nerves (Fig. 4A), the ring gland (Fig. 4B), the brain lobes (Fig. 4C), the AMP clusters in the larval gut (Fig. 4D), and some cells in the brain stem (Fig. 4A and B). Interestingly, a cluster of brain cells with higher staining levels co-stained with an anti-Dilp2 antibody, identifying these cells as Insulin <u>p</u>roducing <u>c</u>ells (IPCs) (Fig. 4E). These cells might be candidates responsible for delaying larval growth and pupation.

**Figure 4:**
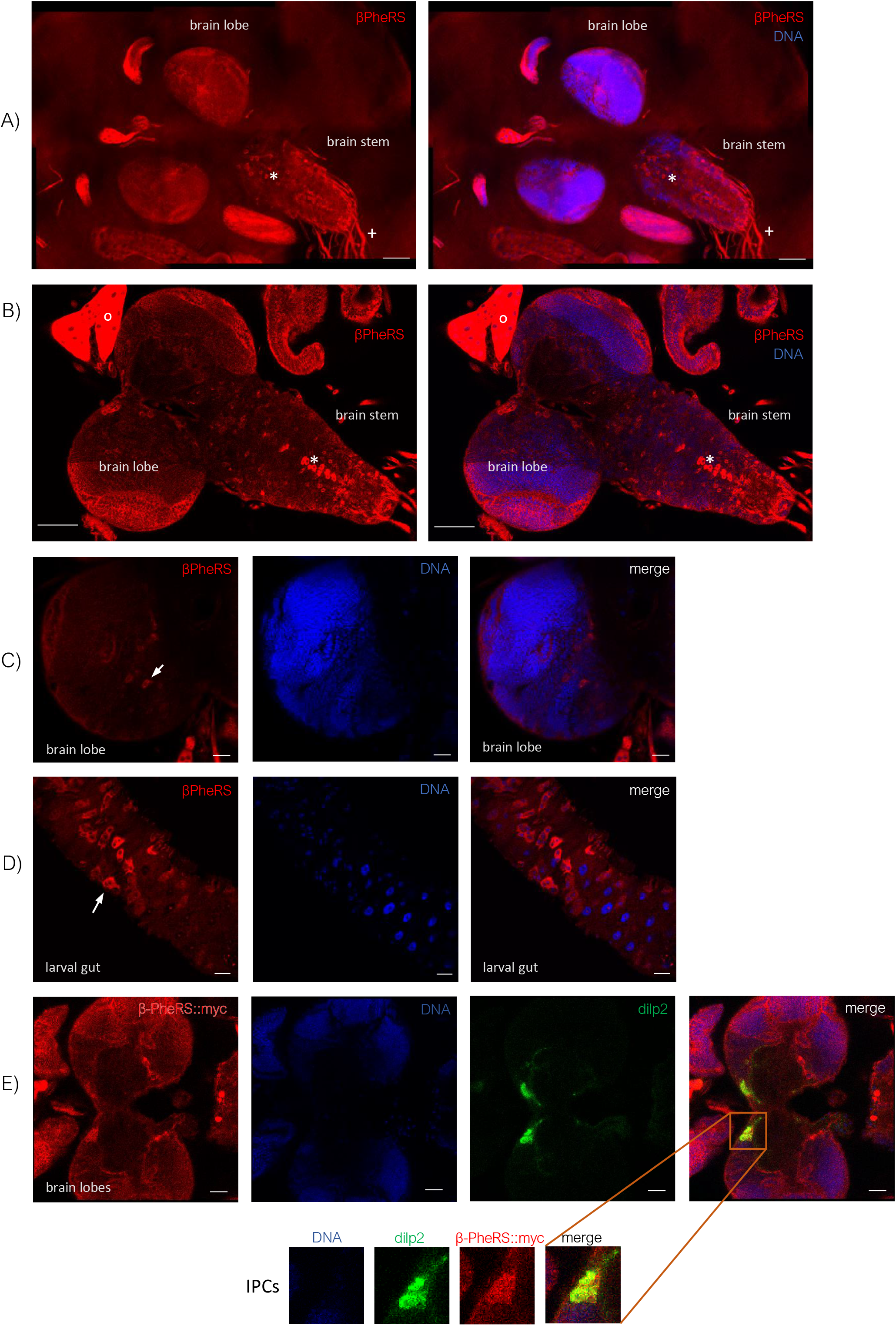
Accumulation pattern of β-PheRS upon α-/β-PheRS overexpression with *tub-Gal4* in larval tissue. In the larval brain, β-PheRS accumulates (A) in the segmentally organized nerves (+), (A) and (B) in some not identified cells in the brain stem (*) and (B) in the ring gland (o). Scale bar is 50 µm. (C) In the brain lobes, single neurons display elevated β-PheRS (arrow). Scale bar is 20 µm. (D) In the gut, high accumulation is seen in the AMP clusters (arrow). Scale bar is 20 µm. (E) In the brain lobes, IPCs display elevated β-PheRS. α-/β-PheRS::myc overexpressing brain lobes stained for Dilp2 and Myc. Dilp2 marks the IPCs. Scale bar is 20 µm.

### Tissue-specific inhibition of overexpression averts the developmental delay

Induction of slow growth and developmental delay appears to require strong overexpression because only a strong driver (*tub-Gal4*) led to the slow growth, while a weaker driver, which is also considered to be a ubiquitous one (*actin-Gal4*) did not induce it. Tissue-specific drivers might also not be strong enough to induce a measurable developmental delay. We, therefore, decided to use the strong, ubiquitous *tub-Gal4* driver in combination with tissue-specific expression of the inhibitor of Gal4, Gal80, to test whether overexpression of PheRS in a specific tissue causes these phenotypes. Suppression of the delay by Gal80 might identify the cells that express Gal80 as candidate cells and tissues where α-/β-PheRS overexpression causes the developmental delay. Two neuronal Gal80 inhibition drivers, the *elav-Gal80* and the *nSyb-Gal80* averted the developmental delay fully or to a high extent, with a remaining 1-day delay for the α-/β-PheRS^B5a^ or α-/β-PheRS^B5b^ overexpression (Fig. 5A and B). The *Su(H)GBE-Gal80* driver is among others reported to be expressed not only in the neuronal tissue but also in the PC cells in the larval gut (Issigonis and Matunis 2010). This driver partially rescued the developmental delay down to 7 days instead of 9-10 days for α-/β-PheRS^B5a^ and α-/β-PheRS^B5b^ overexpression. Furthermore, it completely averted the developmental delay induced by α-/β-PheRS^+^ overexpression (Fig. 5C and Table 3). In contrast, the control *eye-Gal80* driver did not avert the developmental delay caused by overexpression of α-/β-PheRS^+^ and α-/β-PheRS^B5b^ but partially averted the developmental delay caused by the overexpression of the mutant α-/β-PheRS^B5a^ (Table 3). This partial effect was not further analyzed in this study.

**Figure 5:**
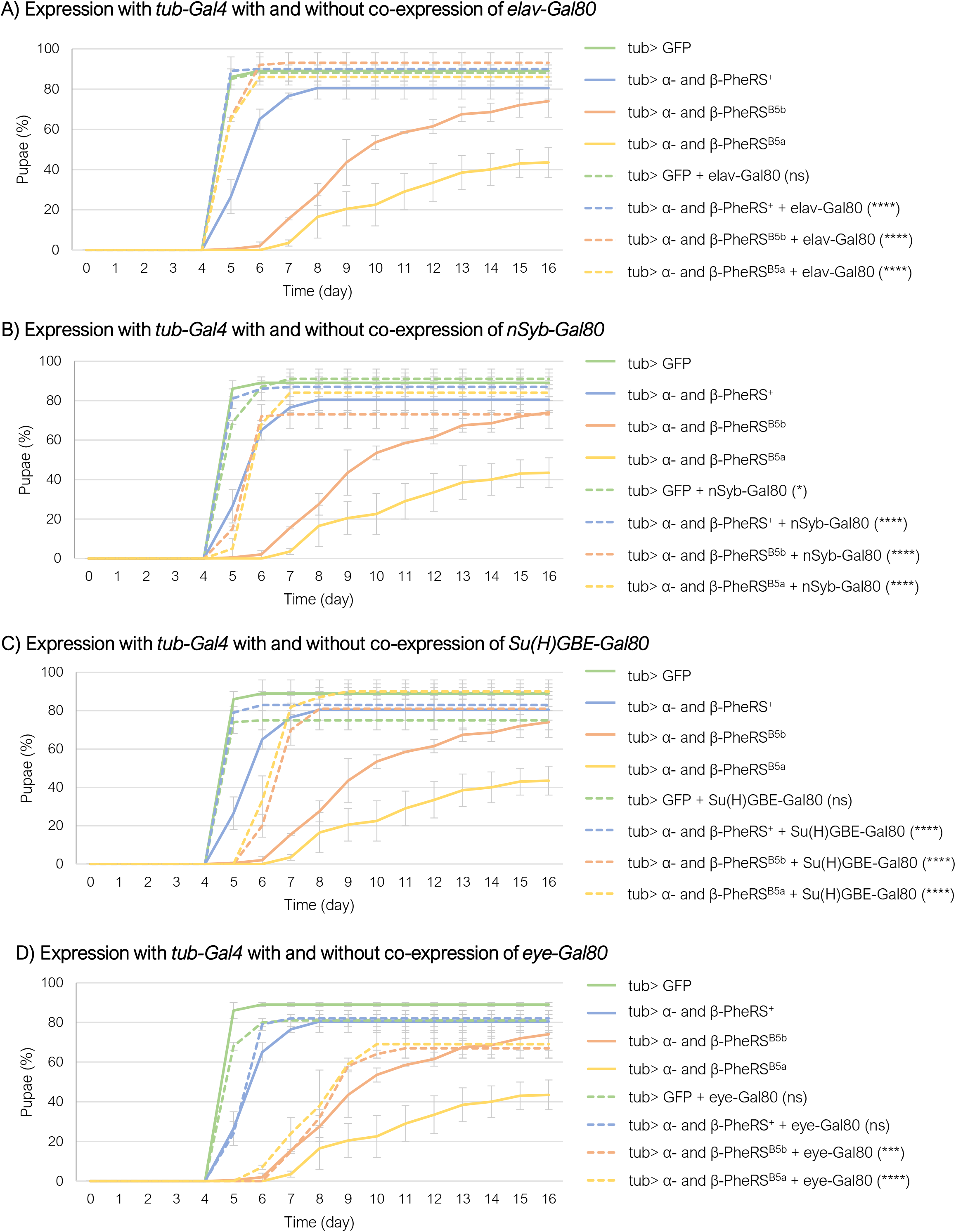
Overexpression of GFP or α-/β-PheRS^X^ with tubulin-Gal4 with or without the addition of a tissue-specific inhibitor of Gal4. (A) elav-Gal80, (B) nSyb-Gal80, (C) Su(H)GBE-Gal80, or (D) eye-Gal80 (Mann-Whitney-U-Test). All experiments were performed in duplicates with 50 larvae each. Graphs represent median ± SD. Mann-Whitney-U-Test was used to compare results to the control. p-value not significant (ns) > 0.05, * ≤ 0.05, ** ≤ 0.01, *** ≤ 0.001, **** ≤0.0001.

The suppression of the developmental delay by neuronal and gut tissue-specific inhibition of overexpression (Table 3, Suppl. Table S1) points to potential roles of the brain and the gut in the α-/β-PheRS^X^ induced developmental delay.

### High expression of α-/β-PheRS^X^ in specific tissues or cell types delays development

Different tissue-specific drivers were also used to narrow down candidate tissue(s) involved in the induction of the developmental delay. Upon driving overexpression of α-/β-PheRS^X^ with these drivers, we assessed the developmental speed of these animals. To facilitate a higher throughput testing, not the time to pupation but the time till the flies eclosed from the pupal case was determined in these experiments. Overexpression of α-/β-PheRS^X^ in the nervous system, a subset of neurons, the optic lobe, the fat body, the salivary glands, ring glands, enterocytes, stem cells, the AMP cells, neurons and glia, the motor neurons and others did not lead to any developmental delay (Table 4). As mentioned before, the interpretation of negative results is difficult because we cannot easily discriminate between a driver that is too weak and one that does not express in the tissue that causes the developmental delay.

**Table 4:** Cell- or Tissue-specific drivers tested for induction of a developmental delay

| Drivers used to overexpress α-β-PheRS<sup>x</sup> | | | Drivers used to overexpress 1x α- and 2x β-PheRS<sup>x</sup> | | |
| --- | --- | --- | --- | --- | --- |
| Focus on Tissue/Cells<br>(can additionally be expressed in other Tissue/Cells) | Gal4 driver | Delay | Focus on Tissue/Cells<br>(can additionally be expressed in other Tissue/Cells) | Gal4 driver | Delay |
| Neuronal | *elav-Gal4* | no | Neuronal | *elav-Gal4* | no |
| Neuronal | *nSyb-Gal4* | no | Neuronal | *nSyb-Gal4* | yes |
| Specific neurons | *5OHO5-Gal4* | no | Specific neurons | *5OHO5-Gal4* | no |
| Optic lobe | *G124<sup>c855a</sup>-Gal4* | no | Optic lobe | *G124<sup>c855a</sup>-Gal4* | no |
| Fat body | *0.68Lsp2-Gal4* | no | Eye | *longGMR-Gal4* | no |
| Eye | *ey-Gal4* | no | Fat body | *0.68Lsp2-Gal4* | no |
| Wing | *wg-Gal4* | no | Neuron and Glia | *nrv2-Gal4* | no |
| Wing | *en-Gal4* | no | Motor neuron | *Hb9-Gal4* | no |
| Fat body | *ppl-Gal4* | no | CCHa2<sup>+</sup> cells | *CCHa2-Gal4* | yes |
| Fat body | *3.1LSP2-Gal4* | no | Pros<sup>+</sup> cells | *pros-Gal4* | yes |
| Salivary gland | *fkh-Gal4* | no |  |  |  |
| Ring gland | *phm-Gal4* | no |  |  |  |
| Enterocytes | *NP1-Gal4* | no |  |  |  |
| Stem cells | *delta-Gal4* | no |  |  |  |
| AMP cells | *esg-Gal4* | no |  |  |  |
| Neurons and Glia | *nrv2-Gal4* | no |  |  |  |
| Motor neurons | *Hb9-Gal4* | no |  |  |  |
| IPCs | *dilp2-Gal4* | no |  |  |  |
| Ubiquitous | *actin-Gal4* | no |  |  |  |
| Ubiquitous | *tub-Gal4* | yes |  |  |  |

Adding to the α-/β-PheRS^+^ overexpression, a second copy of *UAST-β-PheRS*^*+*^ increased the severity of the phenotype drastically (Fig. 2). Using this effect, some Gal4 drivers were also tested by overexpressing one copy of *α-PheRS* with two copies of *β-PheRS*^*X*^ and the time till the flies hatched was determined. However, *UAST-α-PheRS* and two copies of *UAST-β-PheRS*^*X*^ driven with a neuronal, a specific neuron, an optic lobe, an eye, a fat body, a neuron and glia cell, and a motor neuron-specific driver did not lead to any developmental delay (Table 4). In contrast, using the neuronal driver *nSyb-Gal4*, a delay in hatching was observed. A time-to-pupation assay was subsequently also performed and a developmental delay of 1 day was measured when this driver was used to overexpress 1x α- and 2x β-PheRS^B5a^ or 1x α- and 2x β-PheRS^B5b^ (Fig. 6A). This positively identifies nSyb-Gal4^+^ cells as cells where overexpression of 1x α- and 2x β-PheRS^B5a^ or 1x α- and 2x β-PheRS^B5b^ elicits a developmental delay.

**Figure 6:**
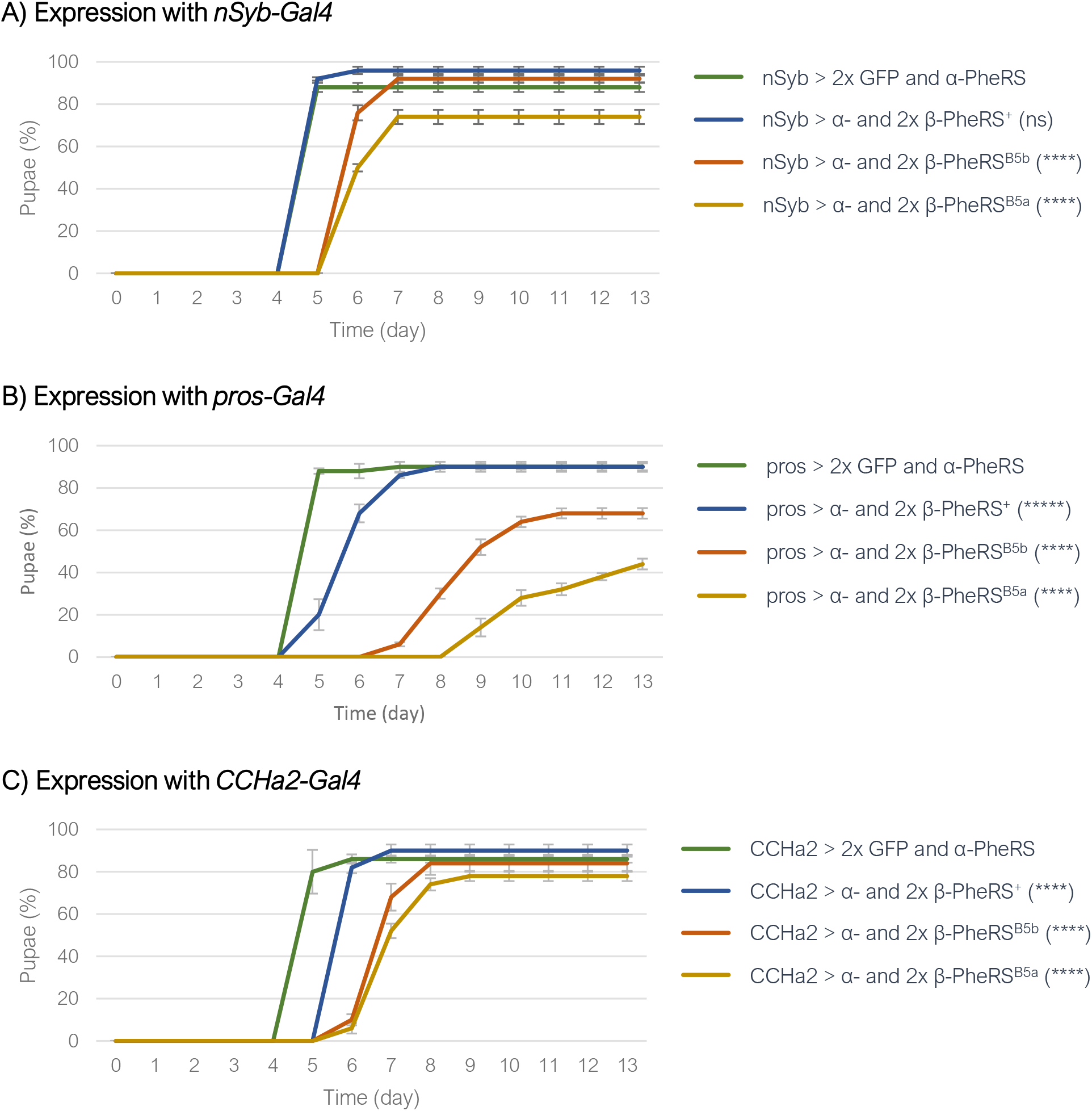
Effects of overexpression of PheRS with different drivers. Time to pupation when control GFP or 1x α- and 2x β-PheRS^X^ were overexpressed with the (A) *nSyb-Gal4* driver, (B) *pros-Gal4* driver, or (C) *CCHa2-Gal4* driver. All experiments were performed in triplicates with 50 larvae each. Graphs represent median ± SD. Mann-Whitney-U-Test was used to compare results to control. p-value not significant (ns) > 0.05, * ≤ 0.05, ** ≤ 0.01, *** ≤ 0.001, **** ≤0.0001.

The *elav-Gal4* did not show a developmental delay while *nSyb-Gal4* did induce one. The neuronal driver *nSyb-Gal4* is a stronger driver than the *elav-Gal4* (Storkebaum et al 2009), but, additionally, the expression patterns of the two drivers differ, too (Suppl. Fig. S2A and S2B). Furthermore, even though Gal4 drivers are often used as tissue-specific drivers, some of them express to some extent in other tissues as well. This was reported already by others (Chen et al 2016, Weaver et al 2020) and observed by us. The neuronal *nSyb-Gal4* driver additionally showed expression in some enteroendocrine (EE) cells throughout the larval gut while *elav-Gal4* showed only expression in EE cells in a small unidentified part of the larval gut (Suppl. Fig. S3A) and no expression in EE cells in the rest of the gut (Suppl. Fig. S3B).

To further test if EE cells in the gut are candidate cells for the induction of the developmental delay, overexpression with *prospero-Gal4* (*pros-Gal4*) was used. In the gut, *pros-Gal4* is an EE cell-specific driver. Overexpression of 1x α- and 2x β-PheRS^+^ with the *pros-Gal4* led to a developmental delay of 1 day and overexpression of 1x α- and 2x β-PheRS^B5a^ or 1x α- and 2x β-PheRS^B5b^ with the same driver to a delay of 4-7 days (Fig. 6B). Over-accumulation of β-PheRS in Pros^+^ cells in the brain was also observed using this overexpression protocol (Fig. 7). This positively identifies at least a subset of the pros-Gal4^+^ cells as cells where overexpression of 1x α- and 2x β-PheRS^+^, 1x α- and 2x β-PheRS^B5a^ or 1x α- and 2x β-PheRS^B5b^ (1x α- and 2x β-PheRS^X^) elicits the prolongation effect. Whether this is mediated by the gut cells or *pros-Gal4* positive cells in another tissue cannot be deduced from these results.

**Figure 7:**
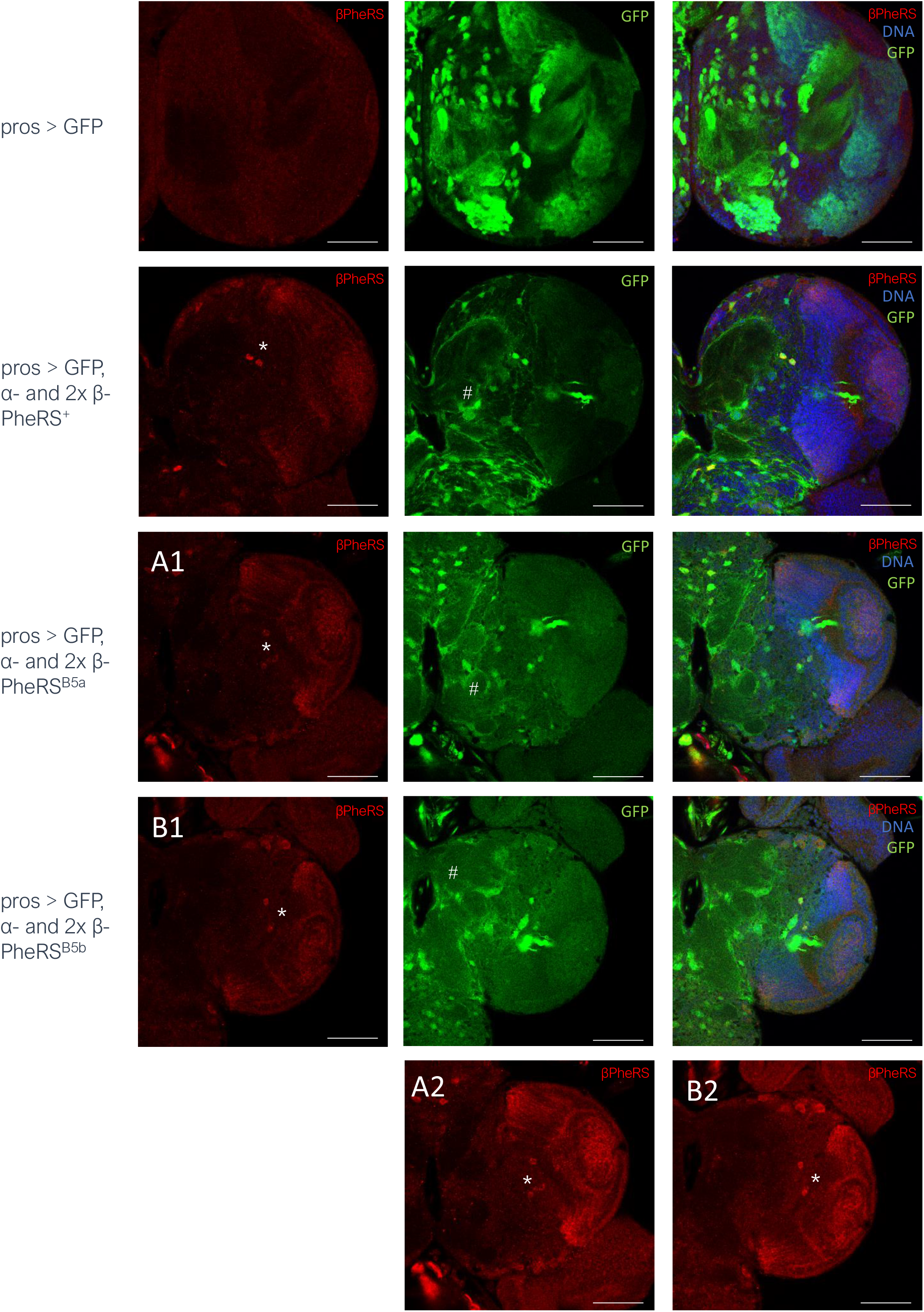
Effect of *Pros-Gal4* driven expression of GFP or 1x α- and 2x β-PheRS^X^. Larval brains were stained for β-PheRS and DNA (Hoechst). β-PheRS accumulates in some GFP^+^ cells (*). The background GFP signal (#) stems from the landing platform used for the PheRS constructs. Scale bar is 50 µm. Pictures were taken with the same settings, except for A2 and B2, which are the same images as A1 and B1, respectively, taken with higher laser intensity. Brain size differs slightly caused by some variation in larval size due to the developmental delay.

### Induction of the developmental delay in CCHa2^+^ cells

Neurosecretory cells and neuropeptides are known to affect the growth rate and feeding behavior in *Drosophila* (Nässel and Zandawala 2019, Nässel und Zandawala 2020). Activation or inhibition of specific subsets of neurosecretory cells as well as reduction or overexpression of neuropeptides can change feeding behavior as well as locomotion activity (Lin et al 2019, Schoofs et al 2014, Zinke et al 1999, Ren et al 2015, Zhan et al 2016, Wu et al 2003).

The IPCs are important regulatory cells for hunger and starvation response. Expression and release of DILP2, 3, and 5, as well as Drosulfakinin (DSK) in these specialized brain cells, regulate feeding and foraging behavior (Lin et al 2019). DILP production in IPCs and their release from the cells are regulated by a variety of upstream factors such as neurosecretory cells and their respective neuropeptides. These neurosecretory cells and the release of their neuropeptides are in turn regulated by other factors sensing nutritional state, food availability, and physiological state of the animal. Many factors involved in regulating hunger and satiety are known, but much remains unknown about the regulation of hunger, satiety, food-seeking, and food intake.

Activation or inhibition of specific neurosecretory cells affects feeding decisions as for example the activation of Hugin-positive neurons induces low food intake and high locomotion activity or roaming (Schoofs et al 2014). In contrast, the activity of Tyrosine hydroxylase (TH)-positive dopaminergic neurons increases feeding, while their inhibition decreases feeding (Bjordal et al 2014). Activation of Taotie neurons, a small cluster of cells in close proximity, but not overlapping the IPCs, induces persistence of hunger signals while inhibition of them induces low feeding (Zhan et al 2016, Lin et al 2019). Furthermore, there are also neuropeptides involved in feeding decisions. Allostatin A (AstA), acts as a satiety signal, inhibiting adult feeding when flies are provided with food (Hentze et al 2015, Lin et al 2019). In contrast, the broad overexpression of Neuropeptide F (NPF) in the nervous system prolongs the feeding phase of third instar larvae (Lin et al 2019), leading to delayed pupation (Wu et al 2003).

β-PheRS might affect some of these specific neurons or neuropeptides, leading to a decreased hunger signal. To investigate whether a cell-autonomous effect of β-PheRS in one of these cells causes the phenotype of a prolonged larval period, we used drivers that are specific for a subset of the neurons with demonstrated functions in satiety and starvation to overexpress α-/β-PheRS. α-/β-PheRS^X^ overexpression with *0098-Gal4, Dh44-Gal4, TH-Gal4, Hugin-Gal4, Dilp2-Gal4, NPF-Gal4, sNPF-Gal4, SIFa-Gal4, Taotie-Gal4, AstA-Gal4* did not lead to any delayed pupation. Furthermore, using an additional copy of *UAST-β-PheRS*^*x*^ with the *TH-Gal4* and *dilp2-Gal4* drivers did also not lead to a delayed pupation. However, the *CCHa2-Gal4* driver led to a very weak delay in pupation when combined with α-/β-PheRS^X^ (one copy of β-PheRS^X^) and to a clear developmental delay when combined with 1x α- and 2x β-PheRS^X^ (Fig. 6C). 1x α- and 2x β-PheRS^+^ overexpression led to a delay of 1 day in pupation onset while 1x α- and 2x β-PheRS^B5a^ or 1x α- and 2x β-PheRS^B5b^ overexpression led to 2 days delay (Table 5). Accumulation of β-PheRS in CCHa2^+^ cells was also observed under these conditions (Fig. 8). This positively identifies *CCHa2-Gal4*^*+*^ cells as cells where overexpression of 1xα- and 2xβ-PheRS^X^ elicits an L3 prolongation effect. We note that overexpression of the β-PheRS^B5a/b^ mutant protein causes less accumulation of the β-PheRS protein signal in CCHa2^+^ cells than the wild type overexpression. We will discuss this result and its implication extensively in the Discussion.

**Table 5:** Median time till larvae pupated when driven with nSyb-, CCHa2- and pros-Gal4

|  | *nSyb-Gal4* | *CCHa2-Gal4* | *pros-Gal4* |
| --- | --- | --- | --- |
| GFP o/e | 5 | 5 | 5 |
| α- and 2x β-PheRS<sup>+</sup> o/e | 5 | 6 | 6 |
| α- and 2x β-PheRS<sup>B5a</sup> o/e | 6 | 7 | 11 |
| α- and 2x β-PheRS<sup>B5b</sup> o/e | 6 | 7 | 9 |

**Figure 8:**
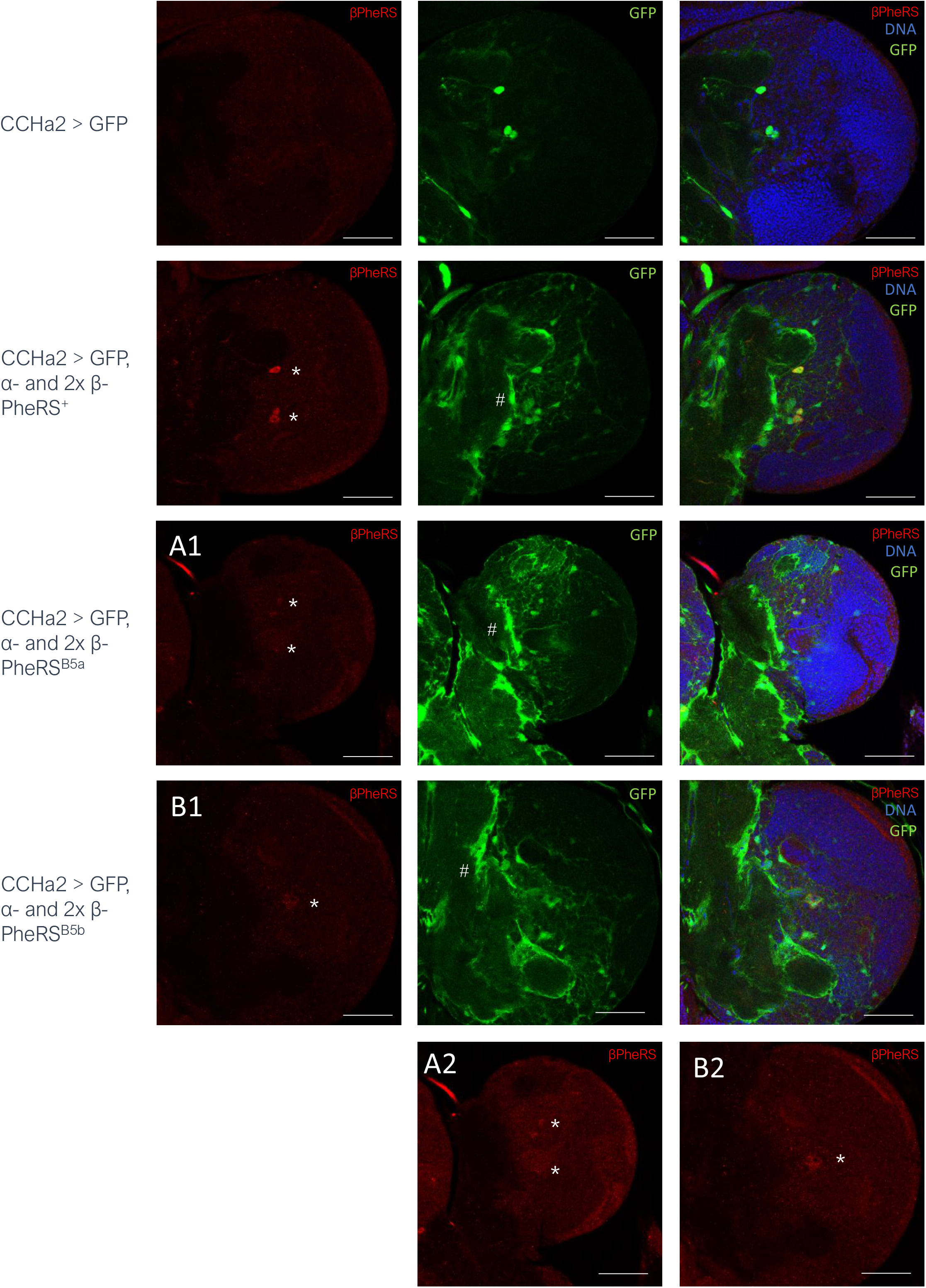
Effect of *CCHa2-Gal4* driven expression of GFP or 1x α- and 2x β-PheRS^X^. Larval brains were stained for β-PheRS and DNA (Hoechst). β-PheRS accumulates in some GFP^+^ cells (*). The background GFP signal (#) stems from the landing platform used for the PheRS constructs. Scale bar is 50 µm. Pictures were taken with the same settings, except for A2 and B2, which are the same images as A1 and B1, respectively, taken with higher laser intensity. For the merged panels of A1 and B1, the red channel images taken with the higher laser power (A2 and B2 images) were used. Brain sizes differ slightly because of the variation in larval size due to their developmental delay.

The *CCHa2-Gal4* driver is expressed in a subset of neurons in the brain (Fig. 8, Suppl. Fig. S2), the gut (Suppl. Fig. S3), and the fat body (not shown, Sano et al 2015). The fat body-specific drivers *ppl-Gal4, 0*.*68Lsp2-Gal4, 3*.*1Lsp2-Gal4* did not lead to any prolongation phenotype. Therefore, we do not expect the fat body to be important for the prolongation phenotype caused by overexpression of *CCHa2-Gal4*, although we cannot rule out that the three fat body drivers express Gal4 at too low levels. The best candidates are therefore the CCHa2^+^ neurons and the CCHa2^+^ intestinal cells.

CCHa2 has been described as an appetite-inducing peptide (Ren et al 2015, Li et al 2013) and CCHa2 mutants show reduced feeding activity and a delay in pupation of approximately 3 days (Ren et al 2015). Overexpression of PheRS gives a similar phenotype as *CCHa2* mutants and overexpression of PheRS specifically in CCHa2^+^ cells (with the *CCHa2-Gal4* driver) led to a developmental delay. To test the hypothesis that PheRS overexpression repressed the appetite by repressing CCHa2, we additionally expressed the appetite inducing CCHa2 neuropeptide in larvae overexpressing also PheRS. In the first attempt, CCHa2 was co-overexpressed with α-/β-PheRS^X^ with *tub-Gal4* (Fig. 9A). In the control experiment, GFP was co-overexpressed. Additional expression of CCHa2 with α-/β-PheRS^+^ led to a slight prolongation of the time till pupation. This was unexpected but can be explained by a negative effect of broad overexpression of CCHa2 in all tissues. This would also explain the significant change of GFP and CCHa2 expression compared to GFP expression alone. Despite this possible negative effect of broad overexpression of CCHa2 with *tub-Gal4*, co-overexpression of CCHa2 with α-/β-PheRS^B5a^ or α-/β-PheRS^B5a^ showed a slight but not significant reduction in the developmental delay (Fig. 9A). To avoid side effects from the broad expression of CCHa2 in all tissues, the *CCHa2-Gal4* and *pros-Gal4* drivers were used next to overexpress 1x α- and 2x β-PheRS^X^ with the addition of a *UAST-CCHa2* construct or a *UAST-GFP* construct as control. The *pros-Gal4* driver slightly rescued the prolongation of the larval phase if CCHa2 was co-overexpressed with α-/β-PheRS^+^ and slightly but not significantly when CCHa2 was co-overexpressed with α-/β-PheRS^B5a^ or α-/β-PheRS^B5b^ (Fig. 9B). A more striking effect was seen with the *CCHa2-Gal4* driver. This led to a clear reduction in time to pupation when CCHa2 was co-overexpressed compared to the control where GFP was co-overexpressed with 1x α- and 2x β-PheRS^X^ (Fig. 9C). CCHa2 was fully able to avert the developmental delay when it was co-expressed. This suggests that the expression of CCHa2 in CCHa2^+^ cells is necessary to avert or override the developmental delay caused by overexpression of PheRS^X^.

**Figure 9:**
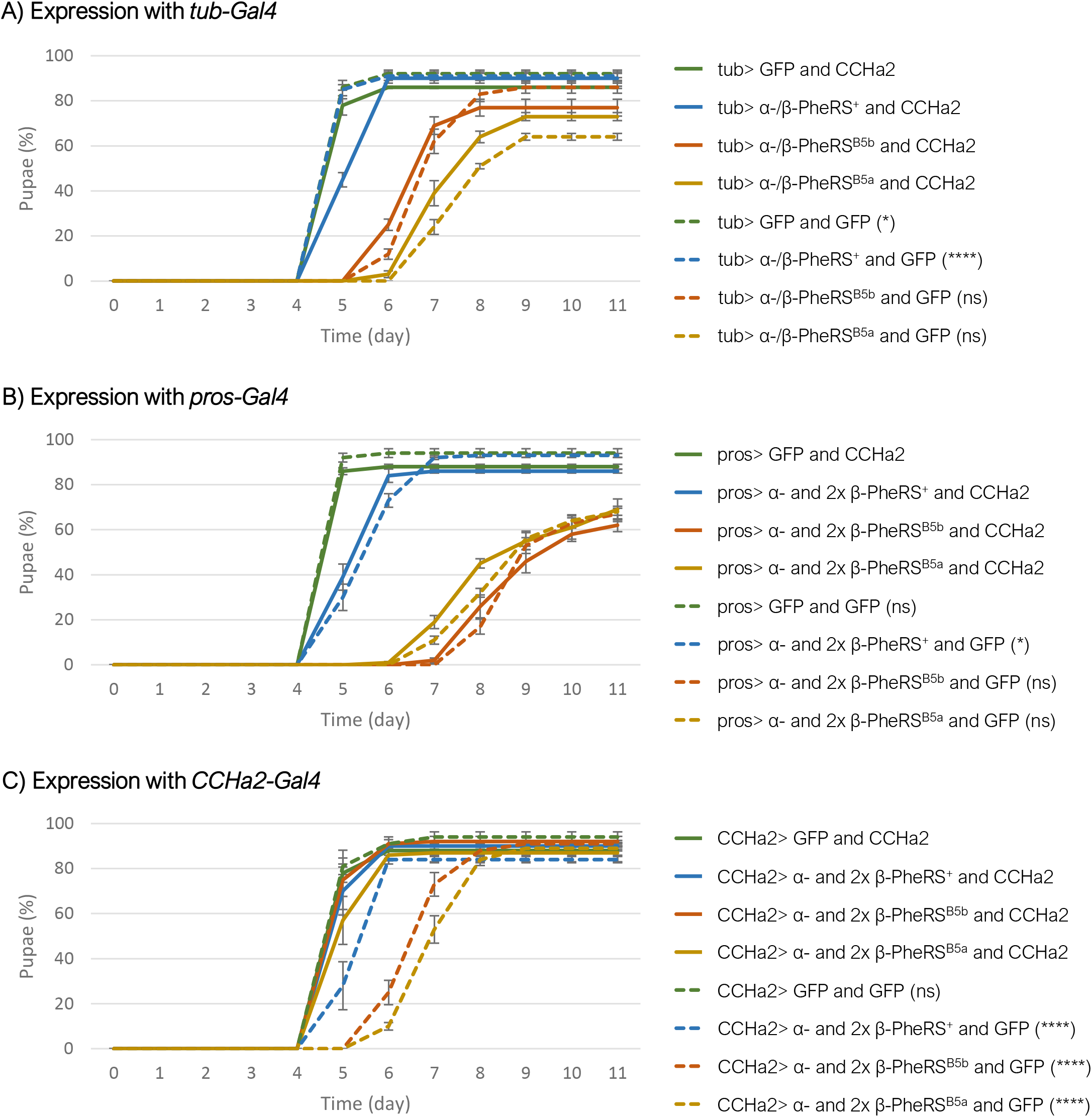
Co-overexpression with CCHa2. (A) *tub-Gal4* driven overexpression of GFP or α-/β-PheRS^X^ with GFP or CCHa2. (B) and (C) Overexpression of GFP or 1x α- and 2x β-PheRS^X^ with GFP or CCHa2. The (B) *pros-Gal4* or (C) *CCHa2-Gal4* drivers were used. All experiments were performed in triplicates with 50 larvae each. Graphs represent median ± SD. Mann-Whitney-U-Test was used to compare CCHa2 to GFP addition. p-value not significant (ns) > 0.05, * ≤ 0.05, ** ≤ 0.01, *** ≤ 0.001, **** ≤0.0001.

### Protein levels upon overexpression of α-/β-PheRS^X^

Mass spectrometry analysis of L1 larvae overexpressing α-/β-PheRS^+^ in an *α-/β-PheRS*^*+*^ background showed an increase in α-PheRS and β-PheRS of 4.2x and 4.3x, respectively, with an α-PheRS/β-PheRS ratio of 1.02 (Table 6). In contrast, overexpression α-/β-PheRS^B5a^ or α-/β-PheRS^B5b^ in an *α-/β-PheRS*^*+*^ background led to an increase in β-PheRS of 2.0x and 2.1x, respectively, and an increase in α-PheRS levels of 2.6x. Therefore, overexpression of α-/β-PheRS^B5a^ or α-/β-PheRS^B5b^ led to a ratio of α-PheRS/β-PheRS^X^ of 1.3 and 1.2, respectively. These results show again that β-PheRS^B5a^ and β-PheRS^B5b^ are less stable, resulting in half the amount of stable protein upon overexpression compared to β-PheRS^+^. Due to the mutual stabilization of the subunits, we would further expect a α-/β-PheRS^X^ ratio of 1. The increased ratio upon overexpression of α-/β-PheRS^B5a^ or α-/β-PheRS^B5b^ possibly results from similar synthesis levels as in α-/β-PheRS^+^, but higher turn-over of the less stable β-PheRS^B5a^ and β-PheRS^B5b^. This result confirms the observation that the CCHa2-driven overexpression leads to lower levels of mutant β-PheRS signal in the CCHa2+ neurons even though the overexpression of the mutant protein causes the stronger phenotypic effect. We will discuss the apparent correlation between the β-PheRS turn-over, the levels of β-PheRS fragments, and the phenotypic effect in the next section.

**Table 6:** Protein levels in L1 larvae upon overexpression of α-/β-PheRS^X^

| Gene | Protein | Protein level<br>change (x) $\alpha/\beta$-<br>PheRS<sup>+</sup> vs control | Protein level<br>change (x) $\alpha/\beta$-<br>PheRS<sup>B5a</sup> vs<br>control | Protein level<br>change (x) $\alpha/\beta$-<br>PheRS<sup>B5b</sup> vs<br>control |
| --- | --- | --- | --- | --- |
| β-PheRS | Q9VCA5 | 4.2x | 2.0x | 2.1x |
| α-PheRS | Q9W3J5 | 4.3x | 2.6x | 2.6x |
| Ratio α-<br>PheRS/β-PheRS | | 1.0 | 1.3 | 1.2 |

## Discussion

### The activity of β-PheRS and α-PheRS in delaying developmental

Mutating two β-PheRS B5 domain residues and motives, respectively, R^353^ (B5a) and GYNN^371-4^ (B5b), respectively, led to lethality, indicating that they are essential or can at least not be replaced by alanine (Table 1). Because the expression levels of β-PheRS^B5a^ and β-PheRS^B5b^ expressed under their endogenous promoter is much lower than the wild-type expression (Table 6; Suppl. Fig. S4) and the resulting phenotype indistinguishable from the *β-PheRS*^*null*^ mutant, the replacement of this residue or motive by alanine causes the degradation of β-PheRS.

To compensate for this instability, we attempted to overexpress *β-PheRS*^*X*^ and, because PheRS subunits are usually only stable if both subunits are overexpressed, we also overexpressed *β-PheRS*^*X*^ together with *α-PheRS*. This co-overexpression, but not the expression of either subunit alone, led to the roaming and developmental delay phenotype, suggesting that the formation of the α-*/*β-PheRS complex is a pre-requisite for the phenotype. Adding an additional copy of *UAS-α-PheRS to the UAS-α-PheRS/UAS-β-PheRS*, rescued the delay while adding a second *UAS-β-PheRS* turned out to extend the delay. This further suggests that the *β-PheRS* isoform produces the delay activity. It might either suggests that *β-PheRS* produces an activity that induces roaming and food avoidance or that it prevents α-PheRS from stimulating feeding and growth.

In all cases tested, overexpression of α-/β-PheRS^B5a/b^ produced a stronger phenotype than α-/β-PheRS^+^ overexpression. This might indicate that the B5 mutations make β-PheRS more active for this secondary activity (and possibly less active for its canonical function). However, there is also evidence for alternative mechanisms. Overexpression of α-/β-PheRS causes only a mild hyperaccumulation in most cell types (Fig. 4; Ho et al., 2022; Lu 2013), because most cells actively control the PheRS levels and cleave and degrade excessive PheRS. Because the B5 mutations cause additional instability of β-PheRS^B5a/b^ (Table 6, Suppl Table S3, Suppl Fig. S3, Suppl Fig. S7), the mutant β-PheRS^B5a/b^ seems to become fragmented even more. The cleavage of the two subunits is accompanied by the formation of stable proteolytic fragments (Ho et al., 2022; Suppl. Fig. S5). It is the more rapid or extensive fragmentation of β-PheRS^B5a/b^ that correlates with the severity of the developmental delay phenotype. This could indicate that a stable proteolytic fragment of β-PheRS might be the causative agent favoring roaming and developmental delay over feeding and growing. In this context, the wild-type R^353^ and GYNN^371-4^ sequences of β-PheRS are thus needed to stabilize β-PheRS and to reduce its fragmentation rate.

The Mass Spectrometry analysis of overexpressing α-/β-PheRS^+^ in larvae showed an approximate 2x higher accumulation of β-PheRS^+^ compared to β-PheRS^B5a/B5b^ despite the presence of the endogenous β-PheRS^+^ in the background (Table 6). Similarly, staining upon overexpression with *CCHa2-* or *pros-Gal4* drivers showed higher accumulation of β-PheRS^+^ than β-PheRS^B5a/B5b^ (Fig. 7 and 8). Despite the lower levels, these α-/β-PheRS^B5a/b^ overexpressing larvae displayed a stronger delay phenotype, suggesting that the delay phenotype might be caused by a higher fragmentation because of their instability. There is precedent for aaRS fragments performing non-canonical activities (Smirnova et al 2012, Park et al 2005 b, Wakasugi and Schimmel 1999, Wakasugi et al 2002, Jung 2015). Particularly relevant for this work is that the α-S fragment of the Drosophila α-PheRS subunit induces a proliferation phenotype and represses Notch activity (Ho et al., 2022).

### Non-cell-autonomous effect of α-/β-PheRS^X^ overexpression

dFOXO overexpression in whole flies suppresses growth and leads to roaming, and overexpression of dFOXO in wings or eyes reduces the size in the respective organ only, demonstrating that dFOXO acts in a cell-autonomous way (Kramer et al 2003). This is not the case for α-/β-PheRS^X^ overexpression. Tissue-specific overexpression in the fat body, the eye- or wing disc did not decrease the size of the fat body, eye, or wing. There was also no change in the morphology of these organs apparent. Importantly, overexpression in all tissues simultaneously, combined with inhibition of expression in neuronal cells and the intestinal EE cells (*elav-Gal80, nSyb-Gal80*) averted most of the prolongation phenotype. These larvae still overexpressed α-/β-PheRS^X^ in most tissues. Nevertheless, they did not show a developmental delay. This points to the neuronal cells and the intestinal EE cells as possible sources of the non-cell-autonomous effect of overexpressed α-/β-PheRS^X^ in reducing growth and extending the larval L3 phase.

### Narrowing down the induction of the developmental delay to a few neuronal and/or gut cells

Overexpression of different combinations of α-/β-PheRS^X^ with *tub-Gal4, nSyb-Gal4, CCHa2-Gal4*, and *pros-Gal4* can lead to a developmental delay. Furthermore, co-expressing the inhibitor of Gal4, Gal80, as *elav-Gal80, nSyb-Gal80*, and *Su(H)-GBE-Gal80* prevented the developmental delay or reduced it at least partially. If only one cell type induces the developmental delay, we expect that Gal4 and Gal80, respectively, are expressed in this cell type in all lines that show an effect. Should more than one cell type be able to induce the phenotype, the evaluation of the result becomes more complex. Starting with the first, simpler assumption, the expression patterns of these drivers led us to focus on the neuronal tissue and the gut (Suppl. Table S1 and S2). *nSyb-Gal4* and *elav-Gal4* were used to express proteins neuronally and the *nSyb-Gal80* and *elav-Gal80* inhibitors were used to inhibit Gal4 drivers neuronally. Recent studies also described the activity of these drivers in the intestinal EE cells (Chen et al 2016, Weaver et al 2020). Chen et al (2016) discovered that *elav-Gal80* inhibits *AstA-Gal4* driven GFP expression in the CNS but also, and unexpectedly, in the EE cells. The same was true for the *nSyb-Gal80*. Chen et al (2016) described EE cell expression of *elav-Gal4* but no EE cell expression of the *nSyb-Gal4* line that they used. Weaver and colleagues described the expression pattern of two different *nSyb-Gal4* drivers and showed EE cell expression for one of the two *nSyb-Gal4* lines. We used the *nSyb-Gal4* line described in that paper as showing no EE cell expression but found that it also drove expression of GFP in the EE cells (Suppl. Fig. S3). From the intensity of the GFP expression signals, it appears likely that the *elav-Gal4* line expressed Gal4 too weakly to induce the developmental delay. The stronger *nSyb-Gal4* line induced a delay, albeit only a weak one. The *CCHa2-Gal4* and *pros-Gal4* lines evidently reached a high enough Gal4 expression level in the important cells to induce the developmental delay. Both drivers show neuronal and gut expression. We have good evidence that the EE cells of the gut are at least not the only cause of the induction of the developmental delay. *Su(H)GBE-Gal80 is* expressed in some neuronal cells and in the PC cells in the larval gut, but not in the intestinal EE cells of the larvae. Despite this, it was able to partially rescue the prolongation phenotype. The fact that the rescue was only a partial one could, indeed, point to some additive effects of the two tissues. Besides the unclear influence of the gut expression, we can conclude that all Gal4 lines that lead to a prolongation phenotype and all Gal80 lines that rescue the prolongation phenotype of *tub-Gal4* overexpression drive expression in the CNS. This strongly indicates an influence of the CNS, more specifically of the CCHa2^+^ and Pros^+^ cells in the CNS, in inducing a prolongation phenotype when α-/β-PheRS^X^ are overexpressed.

### Involvement of hunger and satiety signaling

IPCs and the ring gland are important tissues for the regulators of growth, maturation, and feeding (Lin et al 2019, Nässel and Zandawala 2020). This makes them candidates for providing a link to the α-/β-PheRS overexpression phenotypes of food avoidance, roaming, and developmental delay. Ubiquitous overexpression of α-/β-PheRS^X^ led to the accumulation of β-PheRS in the IPCs and the ring gland. We do not know whether this signal reflects higher levels of the tetrameric α-/β-PheRS or a stable β-PheRS fragment that is still recognized by the antibody. Testing for the effect of overexpressing α-/β-PheRS^X^ only in the IPCs with the *dilp2-Gal4* or in the ring gland with *phm-Gal4* did not lead to any prolongation phenotype. Although not conclusive, this is consistent with the notion that high levels of β-PheRS signal do not necessarily identify cells that produce the non-canonical β-PheRS signal. This observation is also consistent with the result that *CCHa2-Gal4* and *pros-Gal4* overexpressing larvae do not show enhanced accumulation of β-PheRS in IPCs even though they show a prolongation phenotype. The same is true for the ring gland accumulation of β-PheRS. Larvae overexpressing α-/β-PheRS^X^ with *tub-Gal4*, but having the expression blocked in the central nervous system by co-expression of *elav-Gal80*, still accumulate high β-PheRS levels in the ring glands (Suppl. Fig. S6) and this does not lead to a developmental delay. β-PheRS accumulation in IPCs or the ring gland is, therefore, very unlikely to be the main cause of the prolongation phenotype.

Overexpression of 1x α- and 2x β-PheRS^X^, with the *CCHa2-Gal4* driver leads to food avoidance and a developmental delay, indicating a possible involvement of hunger and satiety regulation in promoting the developmental delay. The CCHa2 neurons act upstream of the IPCs and are considered to have a role in linking food availability to growth (Lin et al 2019, Sano et al 2015, Sano 2015). Overexpressing α-/β-PheRS in these neurons, therefore, appears to be the most likely mechanism of extending the larval phase. We would then expect that PheRS overexpression in CCHa2 neurons acts on their signaling to the IPC cells in a manner that induces a food avoidance or a roaming phenotype. The additional expression of the appetite-inducing peptide CCHa2 to these 1x α- and 2x β-PheRS^X^ overexpressing larvae (with the CCHa2-Gal4 driver) averted the developmental delay. This is consistent with a possibly appetite reducing effect of α-/β-PheRS^X^ overexpression and a possible activity of α-/β-PheRS in modulating the CCHa2 signal. The mechanism how β-PheRS modulates the CCHa2 signal still needs to be discovered. The present evidence points to a β-PheRS fragment as the best candidate activity (Suppl. Fig. S5 and Suppl. Fig. S7). Increasing production of β-PheRS fragments by different means (overexpression, destabilizing mutations) correlates with food avoidance and reduced growth. Because this phenotype is rescued by co-overexpression of CCHa2 in the same cells, it appears that high levels of one or more β-PheRS fragments might inhibit CCHa2 expression or its activity on a common downstream target. The significance of our findings may reach beyond Drosophila research. Fragmentation of PheRS (FARS) has also been observed in humans (Greene et al, 2015 a and b) and mutations in human *β-PheRS*, the *FARSB* gene, can lead to problems in gaining weight (Xu et al 2018, Antonellis et al 2018, Zadjali et al 2018). These mutations cause lower levels of FARSB to accumulate, too, indicating that the mutant FARSB is also destabilized. Because of the resemblance of the PheRS and FARS protein behavior and growth problems of the destabilizing mutants in flies and man, the fly result presented here point to a potential mechanism for the human condition and to possible novel approaches to research ways to correct the balance between hunger and satiety signals in the context of obesity. Nevertheless, further studies on the mechanisms by which *β-PheRS* induces the behavioral effect (food avoidance) and the reduced weight gain (developmental delay) are still needed and the Drosophila research could again be useful for such studies.

## Acknowledgments

We thank Jiongming Lu for the initiation of these project. He researched and planed the 5 β-PheRS B5 domain mutations. We also thank him for the sequence alignment in Fig. 1 that identified the conservation of the different B5 domain residues. We thank Barbara Sele for the construction of the B5 domain mutants, Thomas Gentinetta for generating the genomic *β-PheRS* construct and Martin Bergert and Anita Walther for generating the antibody against β-PheRS and identifying and describing the a1103 *β-PheRS*^*null*^ mutation.

Many thanks to Hiroko Sano for giving us the UAST-CCHa2 construct, Alex Gould for the elav-Gal80 line, Hugo Stocker for the nSyb-Gal80 line and I. Miguel-Aliaga for the Su(H)GBE-Gal80 line. We thank Boris Egger, Raffael Koch, Hugo Stocker and Soumya Banerjee for further Gal4 lines and Albena Jordanova for the UAST-GARS line. The Bloomington Stock Center provides invaluable support by distributing many fly lines used here and the DSHB provided multiple antibodies. We also thank Hugo Stocker for the anti-Dilp2 antibody and the PMSCF of the University of Bern for their Mass Spec Analysis.

## Supplementary

### Supplementary Text for Tables S1 and S2

Gal80 inhibitor lines and their reported expression in larvae (Issigonis and Matunis 2010, Chen et al 2016, Weaver et al 2020). Red marked inhibitors suppressed the pupation delay if used to inhibit the effect of tubulin-Gal4 overexpression of α-/β-PheRS^X^. The listed Gal80 lines are expressed in the brain and the gut, even though their cell type specific expression may differ (Suppl. Table S1).

Drivers and their expression patterns described in the literature (Sano et al 2015) and tested in this study with UAS-GFP expression. Green highlighted drivers led to a pupation delay if used to overexpress α-/β-PheRS^X^ (tubulin-Gal4), 1x α- and 2x β-PheRS^X^ (nSyb-, CCHa2- or prospero-Gal4). Not highlighted elav-Gal4 did not induce a pupation delay (Suppl. Table S2). All drivers showed expression in the brain and the gut.

## Legends Supplementary Figures

**Supplementary Figure S1:** Weight development of mixed sex larvae. Control larvae (o/e GFP) and larvae overexpressing α-/β-PheRS^X^. Measurement was started on day 3 after egg lay. Error bares show standard deviation. Graphs represent median ± SD, n = 20. Mann-Whitney-U-Test was used to compare results to control. p-value not significant (ns) > 0.05, * ≤ 0.05, ** ≤ 0.01, *** ≤ 0.001, **** ≤0.0001.

**Supplementary Figure S2:** Expression of Gal4 drivers in the larval brain revealed by UAS-GFP. A) *elav-Gal4*, B) *nSyb-Gal4*, C) *CCHa2-Gal4*, and D) *pros-Gal4*. Scale bar is 50 µm.

**Supplementary Figure S3:** Expression of Gal4 drivers in the larval gut revealed by UAS-GFP. A-B) *elav-Gal4*, C) *nSyb-Gal4*, D) *CCHa2-Gal4* and E) *pros-Gal4*. Scale bar is 20 µm.

**Supplementary Figure S4:** β-PheRS level in *β-PheRS*^*null*^ and rescue larvae. The *β-PheRS*^*null*^ (null) compared to hemizygous *β-PheRS*^*+*^ (con) is shown with or without the addition of a genomic β-PheRS^X^ construct. The *β-PheRS*^*null*^ larvae contain no visible β-PheRS anymore. The same is true for null larvae that also express the genomic construct *β-PheRS*^*B5a*^ and *β-PheRS*^*B5b*^. Note that whereas the B5b result seems clear, there could have been a transfer problem for B5a. However, the reduced accumulation of the B5a and b mutant proteins was also observed by Mass Spectrometry and the mutant proteins have very similar levels (Table 6).

**Supplementary Figure S5:** Western blot of 1x α, 2x β-PheRS^+^, 2x α- and 1x β-PheRS, 1x α- and 1x β-PheRS, and GFP overexpressing larvae stained for β-PheRS and α-PheRS. Marked (*) is the full-size protein. Marked with (°) are two β-PheRS fragments (20 kDa and 45 kDa) formed upon 1x α, 2x β-PheRS^+^ overexpression, and marked with (•) are the two known α-PheRS fragments. The anti β-PheRS antibody was raised and purified against a peptide from the β-PheRS B6 domain.

**Supplementary Figure S6:** Change of β-PheRS levels (red) induced by overexpression with the *tub-Gal4* driver, partially inhibited by co-expression of Gal80. A) Accumulation pattern of β-PheRS upon overexpression of α-/β-PheRS^+^ with *tub-Gal4*. B)-C) Accumulation pattern of β-PheRS upon overexpression of α-/β-PheRS^+^ with *tub-Gal4* with co-expression of the Gal4 inhibitor Gal80 using B) *elav-Gal80* and C) *Su(H)GBE-Gal80*. Accumulation in the ring gland (+), in the IPCs (*), some other neurons (°), and in some cells in the brain stem (#) is seen with tubulin-Gal4 overexpression. Upon inhibition with *elav-Gal80*, the brain did not show any increased accumulation of β-PheRS. Upon inhibition with *Su(H)GBE-Gal80*, the IPCs in the brain lobe and the cells in the brain stem still accumulated high levels of β-PheRS while the brain lobe (besides the IPCs) did not show an increased accumulation of β-PheRS. Scale bar is 50 µm.

**Supplementary Figure S7:** Western Blot of food dependent changes of β-PheRS. Control and α-/β-PheRS^+^ overexpressing larvae were raised on yeast or standard food. Larvae raised on a yeast diet showed an accumulation of an approximately 20 kDa β-PheRS fragment and a decrease of full size β-PheRS compared to larvae raised on standard food. In α-/β-PheRS^+^ overexpressing larvae, starvation reduced this 20 kDa fragment to a lower level than in larvae raised on standard food (or yeast). In α-/β-PheRS^+^ overexpressing larvae, a second fragment of approximately 45 kDa showed a similar abundance pattern in terms of food dependance and starvation as the 20 kDa fragment.

